# Redox Control of S-sulfocysteine Formation in Adenosine Phosphosulfate Reductase

**DOI:** 10.64898/2026.09.04.749436

**Authors:** Matilda Ymeraj, Carlos A. Ramos-Guzmán, Marko Hanževački, Gian Marco Elisi, Giovanni Bottegoni, Adrian J. Mulholland

**Affiliations:** Department of Biomolecular Sciences, University of Urbino, Via Ca’ Le Suore 2/4, 61029 Urbino, Italy; Centre for Computational Chemistry, School of Chemistry, University of Bristol, BS8 1TS, Bristol, United Kingdom; School of Pharmacy, University of Birmingham, Edgbaston, B15 2TT, Birmingham, United Kingdom

## Abstract

Sulfur assimilation fuels bacterial growth by supplying the reduced sulfur required for the biosynthesis of sulfur-containing biomolecules. Adenosine 5′-phosphosulfate reductase (APSR) catalyzes the first reductive step of this pathway, converting adenosine 5′-phosphosulfate (APS) to adenosine monophosphate. This reaction proceeds through nucleophilic attack by catalytic C256, located in the flexible C-terminal tail, on the sulfur atom of APS, forming a thiosulfonate enzyme intermediate. Here, we investigate this reaction in APSR from *Pseudomonas aeruginosa*, an opportunistic pathogen associated with severe infections, particularly in patients with cystic fibrosis. APSR contains an iron-sulfur [4Fe–4S] cluster, which participates in redox steps of the reaction. Here, we show that the redox state of the iron-sulfur cluster also controls the catalytic step. Multiscale molecular simulations investigate how oxidized and reduced cluster states affect APS binding, active site organization, and the nucleophilic attack step. Molecular dynamics (MD) simulations show that the oxidized [4Fe–4S]²⁺ cluster stabilizes substrate interactions and the conformation of the C-terminus, facilitating a catalytically productive orientation of C256. The activation barrier of 17.7 ± 1.7 kcal mol⁻¹ from quantum mechanics/molecular mechanics (QM/MM) umbrella sampling MD simulations at the B3LYP-D3(BJ)/6-31G(d) level of theory is in good agreement with the experimental kinetics. The redox state of the iron-sulfur cluster shows its role in modulating the conformation of conserved K144, which is important for transition state stabilization in the nucleophilic attack. These findings illuminate the mechanism of this *P*. *aeruginosa* target and provide broader insight into the roles of iron-sulfur clusters in controlling enzyme reactivity.

## INTRODUCTION

The increasing spread of multidrug-resistant pathogens has intensified concern particularly over the ESKAPE group (*E. faecium, S. aureus, K. pneumoniae, A. baumannii, P. aeruginosa,* and *Enterobacter* spp.), a leading cause of nosocomial infections and a major public-health threat.^1^ *P. aeruginosa* is a clinically important member of this group, exhibiting broad antimicrobial resistance with biofilm-mediated tolerance that limits the effectiveness of standard therapies.^2^ This opportunistic Gram-negative bacterium is responsible for a wide range of infections and poses a particular risk in immunocompromised hosts and patients with cystic fibrosis.^3^ The clinical persistence of *P. aeruginosa* in lungs of cystic fibrosis patients is driven by its capacity to adapt to inorganic sulfate limitation in airway mucus by activating a sulfate starvation-induced response. This metabolic reprogramming promotes scavenging of host organic sulfur pools, supporting long-term survival and colonization of the respiratory tract.^4^

The production of sulfur-containing primary metabolites, such as cysteine and methionine, represents a fundamental and highly conserved process across bacteria, plants and fungi, sustaining vital functions even under conditions of extreme nutritional stress.^5^ Central to this are sulfonucleotide reductases (SRs), a diverse family of enzymes that catalyze the first committed step in reductive sulfur assimilation for *de novo* cysteine biosynthesis. This enzyme family comprises two subclasses. The first subclass is adenosine-5′-phosphosulfate (APS) reductase, which harbors a [4Fe–4S] cluster^6^ and is expressed in plants and many bacteria, including *P. aeruginosa* and *Mycobacterium tuberculosis*. The second class is 3’-phosphoadenosine 5’-phosphosulfate (PAPS) reductase, which lacks the cluster and is common in enteric bacteria and yeast (Scheme 1).^7^ Although APS and PAPS reductase exhibit modest sequence conservation with 25-33% pairwise identity and 42% similarity, structural analyses reveal a conserved Rossmann-like fold. This architecture is shared across available crystal structures,^8,9^ including *P. aeruginosa* APS reductase.

Results from biochemical and biophysical investigations of sulfonucleotide reductase activity in both APS and PAPS enzymes are consistent with a two-step ping-pong mechanism^10^ in which the sulfonucleotide undergoes nucleophilic attack by a conserved cysteine present in the flexible C-terminal region of the enzyme to form an enzyme S-sulfocysteine intermediate (Enz-Cys-Sγ-SO_3_⁻). In the subsequent step, the intermediate is cleaved by thioredoxin (Trx), releasing sulfite and regenerating the catalytic thiol (Scheme 2). Hence, despite their differences in substrate specificity, APS and PAPS reductases share the same mechanism for the reduction of sulfonucleotides.^11^

**Scheme 1.**
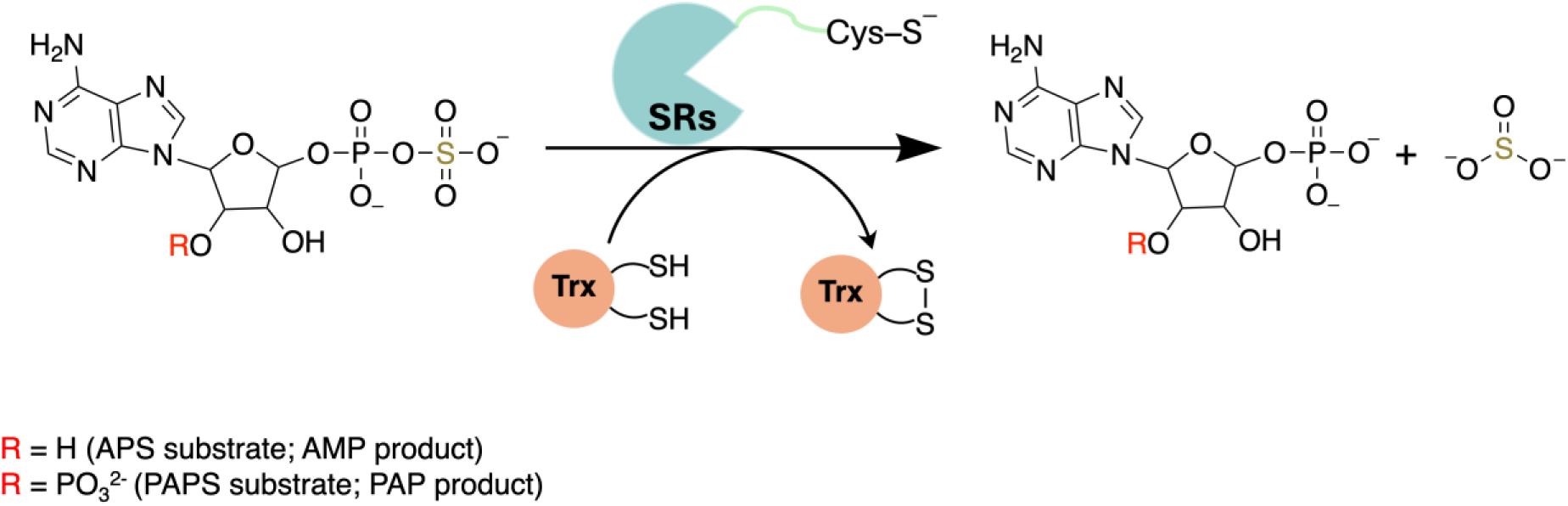
Reaction catalyzed by sulfonucleotide reductases (SRs). Adenosine-5′-phosphosulfate (APS) and 3’-phosphoadenosine 5’-phosphosulfate (PAPS) are converted to adenosine monophosphate (AMP) and adenosine 3’,5’-bisphosphate (PAP), via reaction catalyzed by APS reductase and PAPS reductase, respectively.

**Scheme 2.**
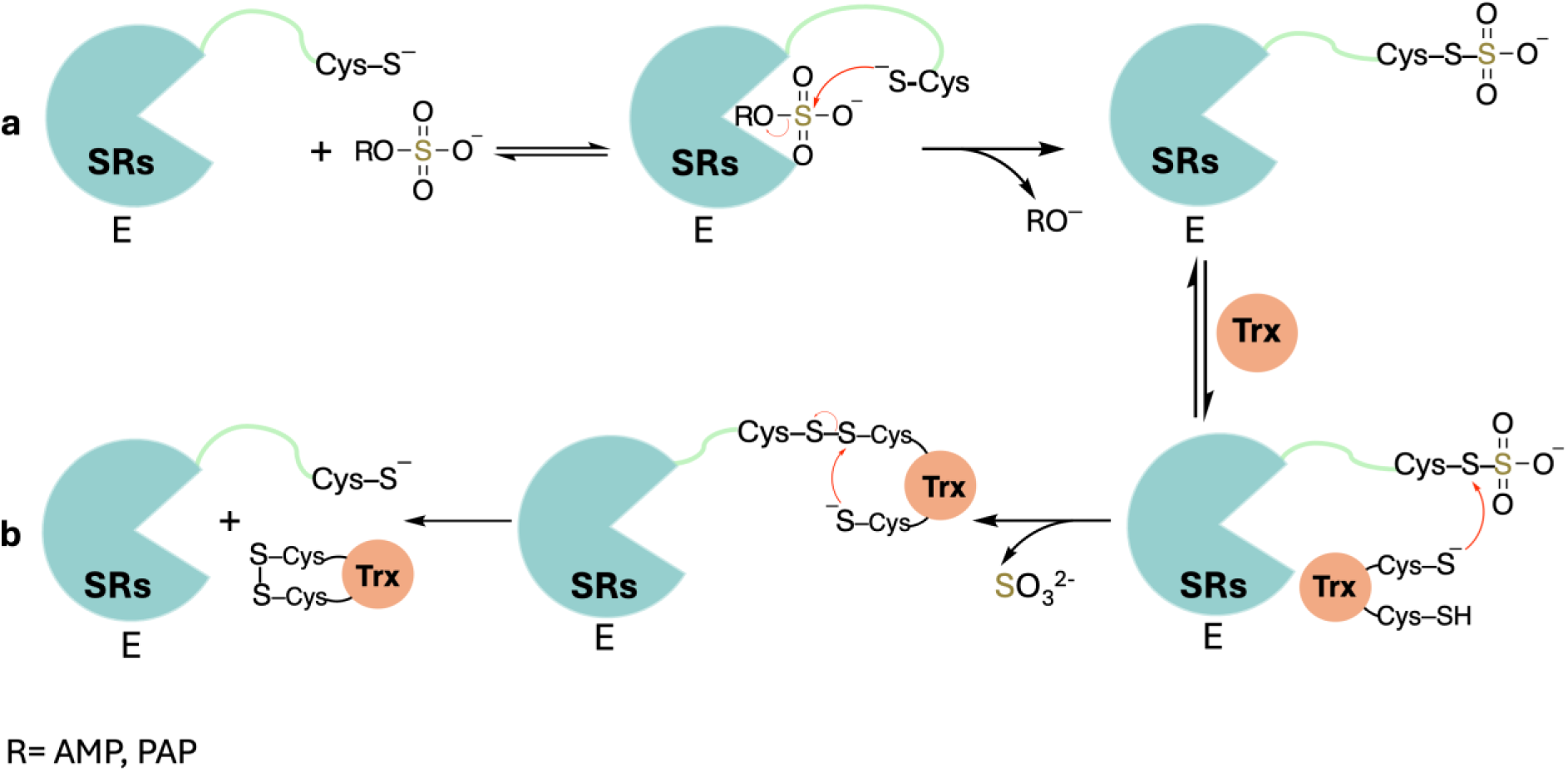
Two-step mechanism of sulfonucleotide reductases (SRs). (a) Step 1. C-terminal Cys-S^−^ attacks the sulfonucleotide to form the covalent Enz-Cys-SγSO_3_ ^−^ intermediate with release of the nucleotide leaving group. (b) Step 2. Thioredoxin (Trx) reduces the covalent enzyme sulfonate, releasing SO_3_ ^2−^ and restoring the catalytic thiolate Enz-Cys-S^−^.

The catalytic cysteine (C256) of APS reductase (APSR) in *P. aeruginosa* is part of the^255^ECG(L/I)H^259^ motif located in a C-terminal tail that also contains a ^242^R(S/E/A)GR(W/F)^246^ substrate-recognition sequence (Figure 1). The latter comprises two conserved arginine residues, namely R242 and R245, that form electrostatic interactions with the phosphosulfate group. Structural and functional analysis suggests a conformational change in which the tail opens for substrate binding, closes to catalyze the reaction and stabilize the thiosulfonate intermediate, and reopens again to release the product (AMP/PAP), enabling Trx-mediated sulfite release and enzyme regeneration.^12,13,14^ In *P. aeruginosa*, the highly flexible region spanning residues 250-267, which contains the catalytic C256, is unresolved in the only available crystal structure (PDB ID 2GOY), consistent with its proposed mobility within the active site.^13^ Site-directed mutagenesis (C256S) abolishes activity and prevents formation of the enzyme-thiosulfonate intermediate in mass spectrometry experiments, confirming the catalytic role of C256.^15^

**Figure 1.**
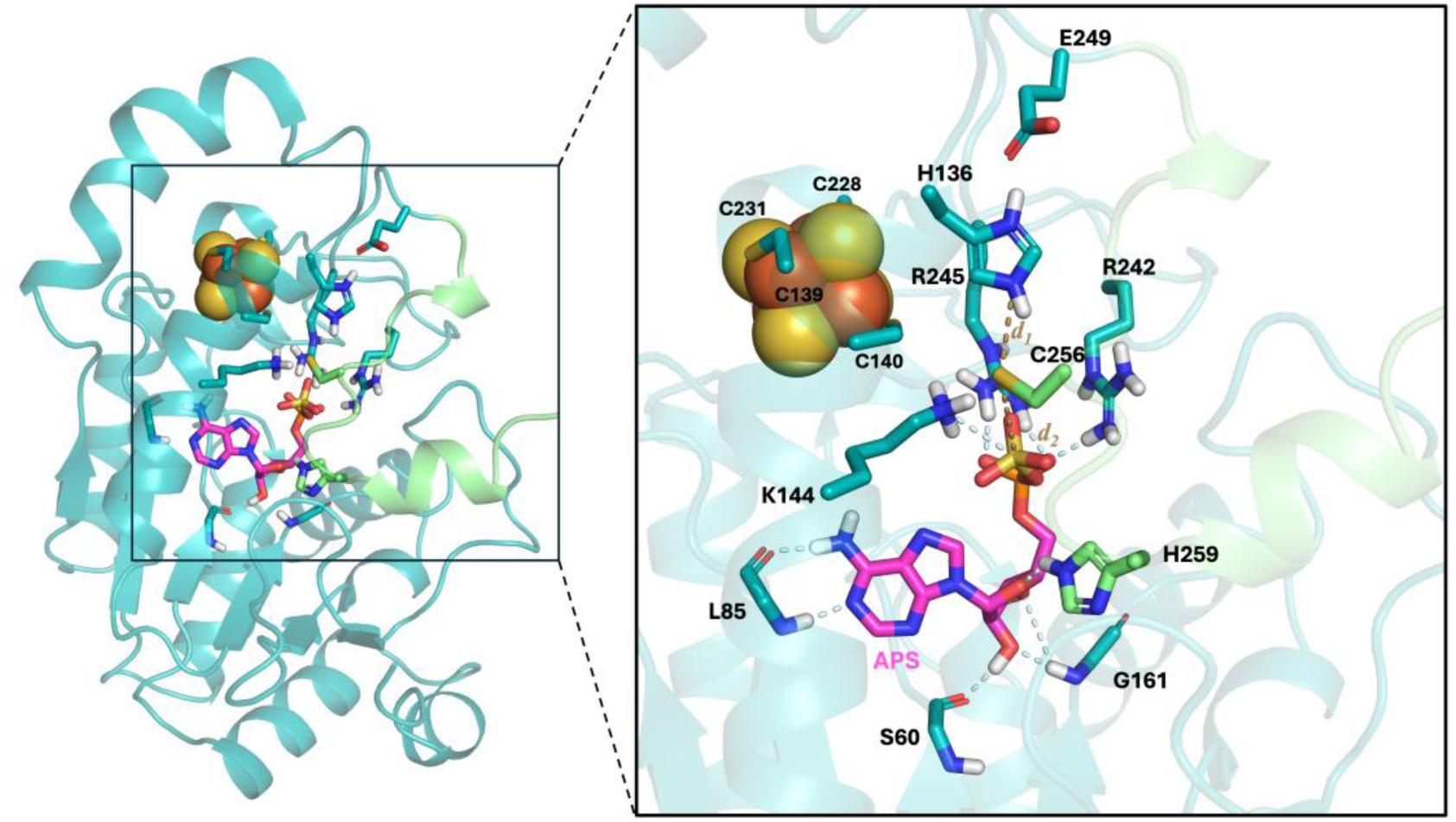
Crystal structure of APSR (PDB ID 2GOY) shown as teal cartoon, with the unresolved C-terminal region (predicted using AlphaFold2 and shown as green cartoon). The bound APS substrate is stabilized via hydrogen bonding of the adenosine moiety with S60, L85, G161, and H259, and through the phosphosulfate group with conserved residues K144, R242, and R245. The predicted C-terminus structure revealed that the sulfur atom of C256 forms a hydrogen bond with H136, which forms a salt bridge with E249, suggesting their potential involvement in a catalytic triad. The iron-sulfur cluster is shown as van der Waals spheres. Catalytically relevant distances (shown as yellow dashed lines) are defined as *d*_1_ = Sγ_C256_-Nε_H136_ and *d*_2_ = Sγ_C256_-S_APS_. Hydrogen bonds are shown as light blue dashed lines. Hydrogens atoms were added using H++ and only polar hydrogens are shown for clarity.

X-ray crystallographic structure of APSR shows that the [4Fe−4S] cluster is coordinated by the conserved ^139^CC-X_80_-CXXC^231^ motif and is positioned approximately 10 Å from the sulfur atom of bound APS, without direct substrate contact. Nevertheless, mutation of the conserved cluster-coordinating cysteines prevented Fe–S cluster incorporation and abolished APSR activity.^16^ Consistently, introducing a [4Fe–4S]-binding site into PAPS reductase increased APS reduction by approximately three orders of magnitude, mainly through acceleration of the chemical step rather than improved substrate affinity.^17^

Mass spectrometry of purified APSR identifies the [4Fe−4S]^2+^ as the predominant oxidation state, with no detectable change across the catalytic states under the tested conditions.^18^ However, this does not rule out transient or thioredoxin-dependent redox changes during regeneration of the active enzyme from the covalent enzyme-sulfonate intermediate. Related radical SAM glycyl radical activating enzymes demonstrate that protein environment and electron delivery partners can enable interconversion between [4Fe–4S]^+^ and [Fe–4S]^2+^ states during single-electron transfer.^19,20^ More broadly, assimilatory sulfate reduction can contribute to intracellular redox homeostasis under sulfide stress by consuming excess reducing equivalents and limiting electron leakage, highlighting the potential importance of redox-sensitive steps within this pathway.^21^ We therefore assess how the [4Fe–4S]^2+^ state and plausible one-electron reduced [4Fe–4S]^+^ state affect APSR active site structure, APS stabilization, and the initial nucleophilic attack step.

In this work, we investigate the first step of the APSR catalytic pathway (Scheme 2) using multiscale molecular simulations. Molecular dynamics (MD) simulations explore the dynamics of bound APS, testing different combinations of protonation states of active site residues H136 and C256. High-level hybrid quantum mechanics/molecular mechanics (QM/MM) MD simulations employing DFT provide an atomistic description of the APSR reaction, supporting a mechanism in which the C256 thiolate performs the nucleophilic attack on APS, generating an enzyme-bound S-sulfocysteine intermediate. The computed activation free energy of 17.7 ± 1.7 kcal mol^−1^ is consistent with the apparent activation free energy of 18.6 kcal mol^−1^, derived at 298.15 K from the experimentally measured catalytic turnover rate (*k*_cat_ = 9.2 ± 0.1 s^−1^).^22,23^ Our simulations demonstrate that the cluster oxidation state controls the organization of the APSR active site; reaction is facilitated in the presence of the oxidized [4Fe-4S]^2+^ cluster. Together, these results reveal the structural and dynamical determinants that govern substrate recognition and reaction in ASPR. This understanding will provide support for design of selective APSR inhibitors and for strategies aimed at modulating microbial sulfur metabolism.

## METHODS

### Modeling of the Michaelis complex and system parametrization

The X-ray crystal structure of *P. aeruginosa* APSR in complex with adenosine 5’-phosphosulfate was obtained from the Protein Data Bank (PDB ID 2GOY)^13^; chain B was processed with the Protein Preparation workflow implemented in the Schrödinger suite.^24^ The unresolved flexible C-terminal region (residues 250–267; see Figure 1 and Figure S1) was built in Prime^25^ employing the corresponding segment of the AlphaFold2^26^ model as a structural template. The incorporated C-terminal region was subsequently refined by OPLS4 minimization,^27^ with the crystallographic structure positionally constrained to preserve the experimental coordinates. Protonation states of all ionizable residues were assigned at physiological conditions using the H++ server.^28^ Standard amino acid residues were represented by the AMBER ff14SB force field.^29^

Geometry optimization of the APS ligand and electrostatic potential (ESP) calculations were performed in Gaussian 16.^30^ The ligand atom-types were assigned using GAFF2^31^ and the atomic charges obtained with restrained electrostatic potential (RESP)^32^ derived at the HF/6-31G(d) level of theory. Metal ions were modelled using the Li-Merz parameter set.^33^ The enzyme-substrate complex was solvated in a cubic TIP3P^34^ water box ensuring a minimum distance of 10 Å between the atoms of the solute and the box edges. Periodic boundary conditions (PBC) were used. Charge neutrality was ensured by adding Na⁺ counterions. All system preparations were performed using AmberTools.^35^

### Iron-sulfur cluster parametrization

Bonded force field parameters for the iron-sulfur [4Fe–4S] cluster were generated using the metal center parameter builder (MCPB.py)^36^ program, employing both small and large model systems. The small model, consisting of the [4Fe–4S] cluster and four coordinating ethane thiolate ligands, was used to derive force constant via the Seminario method^37^ based on vibrational frequency calculations. The large model, comprising the full cluster site with the coordinating cysteine residues capped by acetyl and N-methyl groups, was used for RESP charge fitting. Both models were optimized at the B3LYP/6-31G(d) level of theory. The oxidized and reduced clusters were parametrized separately as [4Fe–4S(Cys)_4_]^2–^ and [4Fe– 4S(Cys)_4_]^3–^, respectively. In the QM calculations, total charges of –2 and –3 were assigned to the oxidized and reduced states, respectively. The spin states of both oxidation states were treated using a broken-symmetry density functional theory (DFT) approach to describe the antiferromagnetic coupling between the iron centers.^38^

### MM molecular dynamics simulations

Molecular dynamics (MD) simulations were used to assess five alternative protonation state configurations of the catalytic C256 residue and the neighboring H136, using the catalytically competent oxidized [4Fe–4S]^2+^ cluster.^39^ The following combinations were investigated using standard AMBER parameters: (i) neutral cysteine (CYS)/neutral histidine *_δ_*-protonated (HID), (ii) CYS/double protonated histidine (HIP), (iii) deprotonated cysteine (CYM)/HIP. Additionally, the custom parameters reported by Pedron et al.^40^ for deprotonated cysteine (CY2) were employed to test the effect of sulfur atom Lennard-Jones in the simulations: (iv) CY2/HIP, and (v) CY2/HID. To further explore the influence of the cluster’s redox state, an additional simulation (vi) was carried out for the CY2/HIP combination using the reduced [4Fe–4S]^+^ cluster; see results for more details.

Systems were subjected to energy minimization consisting of 30,000 cycles, with the initial 500 cycles using the steepest descent algorithm, followed by the conjugate gradient method for the remaining steps, until the root mean square of the gradient plateaued at 3×10⁻³ kcal·mol⁻¹·Å⁻¹. Systems were then gradually heated from 40 K to 300 K over 140 ps using Langevin dynamics.^41^ Equilibration was carried out under NPT conditions at 300 K and 1 atm, employing a Berendsen barostat.^42^ Backbone atoms were restrained during equilibration, with positional restraint force constants gradually decreased from 15 to 0 kcal mol^−1^ Å^−2^ in decrements of 3 units every 1.25 ns. After 6.25 ns, all restraints were fully released, and the systems underwent an additional 1.25 ns of unrestrained equilibration. Production MD simulations were subsequently performed in the NVT ensemble at 300 K for 600 ns per replica, with three independent replicas per system (totaling 1.8 μs per protonation state configuration). All molecular mechanics/molecular dynamics simulations (MM MD) were performed with the AMBER24 suite.^43^ Production MD runs used pmemd.cuda,^44^ with the SHAKE algorithm,^45^ thereby allowing a 2 fs integration timestep. A 10.0 Å cutoff was used to evaluate pairwise nonbonded interactions, and those beyond this value were treated using the particle mesh Ewald (PME) method.^35,46^

Resulting MM MD trajectories were analyzed to construct the free energy surface (FES),^47^ presented in this study for the alternative protonation state configurations of the APSR-substrate complex mentioned above. Both oxidized and reduced cluster states were investigated by QM/MM MD simulations using the CY2/HIP combination. Representative low free energy snapshots from the FESs of the APSR-substrate complex were extracted from the MD trajectories using cpptraj in AMBER24,^35^ and used as starting structures for subsequent QM/MM MD.

### QM/MM molecular dynamics simulations

QM/MM MD simulations were carried out using two different QM size regions (Figure S2). The smaller region (QM1) comprised the phosphosulfate moiety of APS along with the C_β_-truncated side chains of H136, K144, and C256 (totaling 44 atoms). The larger region (QM2) encompassed all atoms from QM1 with the addition of the entire APS substrate, the E249 side chain, the peptide bond between C256 and E255, and four water molecules, yielding a total of 98 atoms. The net charges of the QM regions were –1 (QM1) and –2 (QM2), with a singlet spin multiplicity in both cases. At the QM/MM boundaries, hydrogen link atoms were included to complete the valence of the atoms in the QM region. Electrostatic embedding was employed,^48^ with real-space electrostatic interactions truncated at 8.0 Å.

Umbrella sampling^49^ QM/MM MD simulations were performed to investigate the nucleophilic attack and subsequent APS S-O bond cleavage. Preliminary calculations were performed at the semiempirical PM6^50^ level to select QM region size and optimize force constant values, ranging from 100 to 400 kcal mol^−1^ Å^−2^, ensuring proper window overlap prior to production sampling at the DFT level. The reaction coordinate describing the nucleophilic attack was defined as a linear combination of the breaking S_APS_-O_APS_ and forming S_C256_-S_APS_ bonds. Umbrella windows were distributed at 0.2 Å intervals along the reaction coordinate. To obtain structures for every window along the reaction coordinate, energy-minimized (500 steepest descent steps followed by conjugate gradient optimization for a total of 5,000 cycles) umbrella sampling simulations were made, followed by a 300K equilibration using Langevin dynamics.^41^ The QM/MM umbrella sampling simulations at the PM6 level were performed with the sander module in AMBER24,^43^ employing a 1 fs timestep, with each umbrella window equilibrated for 10 ps and followed by 20 ps sampling production runs. The DFT QM/MM MD simulations were performed using the B3LYP/6-31G(d) level with D3 dispersion corrections and Becke-Johnson (BJ) damping.^51,52^ To reduce the computational cost associated with the larger QM2 region, structures previously equilibrated at the PM6/MM level were used as starting geometries for subsequent 10 ps DFT QM/MM production runs. All DFT QM/MM calculations were executed using GPU-accelerated QUICK^53,54^ interfaced with AMBER24. During umbrella sampling simulations, a harmonic restraint was applied to the H136 Nε-H bond to isolate the energetic contribution of the nucleophilic attack from that of the proton transfer between C256 and H136. The potentials of mean force (PMFs) were obtained using the weighted histogram analysis method (WHAM),^55,56^ with associated errors estimated through block averaging analysis.

Additionally, extensive unrestrained PM6 QM/MM MD simulations were carried out for APS (reactant) and AMP (product) states, considering both the reduced and oxidized [4Fe–4S] cluster, to assess ligand dynamics and active site flexibility. These simulations were conducted for 100 ps per replica (three replicas per system yielding a total of 300 ps), starting from equilibrated structures obtained from the umbrella sampling PM6 QM/MM MD runs.

## RESULTS AND DISCUSSION

### Inspection of AlphaFold C-terminus structure through comparison of APS with PAPS reductase

The inclusion of the predicted C-terminus in the APSR structure (shown as green cartoon in Figure 1) reveals a more complete active site architecture where C256 lies close to H136, while H136 forms a hydrogen bond with E249, suggesting the presence of a catalytic triad. In this conformation, C256 is positioned near the APS sulfate group in a geometry favorable for the nucleophilic attack. Moreover, similarly to PAPS reductase, H259 stabilized APS recognition through a hydrogen bond with the ribose oxygen (Figure 1). This agrees with mutagenesis data that show how mutation of H259 to alanine reduced binding affinity without changing the rate-limiting chemical steps of the reaction.^12^

The conserved ^255^ECG(L/I)H^259^ motif shows a mean pLDDT score of 69.92 ± 3.81, supporting a reliable prediction of the local backbone arrangement in this functionally relevant C-terminal region. To further assess the structural plausibility of the predicted conformation, a Cα alignment was performed between APSR, including the predicted C-terminus, and the experimentally resolved PAPS reductase complex (PDB ID 2OQ2),^14^ as shown in Figure S1. Despite their relatively low sequence identity (25.9%) and similarity (41.6%), the two enzymes share a conserved α/β/α sandwich core and exhibit similar active site architectures, as reflected by a Cα root-mean-square deviation (RMSD) of ∼1.0 Å. These observations support the plausibility of the C-terminal loop model in APSR; nevertheless, extensive MD simulations were performed to further assess its conformational stability and dynamics depending on the protonation states within the active site.

### Protonation states and stability of the active site in MM MD simulations

p*K*ₐ estimates obtained with H++ yielded values of 6.5 for C256, 7.01 for H136, and >12 for K144. These values indicate that, at physiological pH, the catalytic C256 also exists as a thiolate, while K144 remains protonated. The predicted p*K*ₐ of H136 suggests that its neutral form predominates at pH 7.4, with the Henderson-Hasselbalch equation giving an approximate 5:2 neutral/protonated ratio. Because the H++ server excludes ligands and metal cofactors from p*K*_a_ calculations, the influence of the [4Fe–4S] cluster was not considered during protonation state assignment. Given the proximity of H136 to this negatively charged cluster, its p*K*_a_ is expected to be increased relative to the calculated value, which would favor the positively charged (doubly protonated) form of the residue. Accordingly, MM MD simulations were performed with systematically varied combinations of protonation states for C256 and H136 to evaluate their impact on substrate conformation and active site dynamics.

In the oxidized [4Fe–4S]^2+^ cluster state, root-mean-square fluctuation (RMSF) analysis showed that deprotonation of C256 decreased the mobility of the C-terminal loop comprising residues 250-267 compared to the neutral state, suggesting that the thiolate form contributes to stabilization of the C-terminal loop across both CYM and CY2 parameter sets (Figure S3). In the neutral CYS/HID configuration, reduced electrostatic stabilization of C256 within the active site promoted greater C-terminal loop flexibility, with the mean RMSF of the catalytic C256 increasing 0.96 ± 0.12 Å in the CY2/HIP system to 1.7 ± 1.1 Å in the neutral CYS/HID system (Figure S3). Among the deprotonated cysteine models, CY2/HIP provided greater ligand stability than CYM/HIP, supporting improved preservation of the substrate binding geometry within the active site (Figure S4).

### Thiolate/imidazolium protonation state configuration

Having established the CY2 parameters as an appropriate description of the catalytic thiolate within the catalytic site, the CY2/HIP system with the [4Fe–4S]^2+^ cluster provided the most stable and catalytically relevant substrate binding arrangement. Simulations performed with the default AMBER thiolate parameters (CYM/HIP) showed gradual substrate displacement, associated with water penetration into the active site and disruption of the hydrogen-bonding network (system iii, Figure S4). Simulations performed with the refined sulfur Lennard-Jones parameters for the thiolate (CY2/HIP),^40^ developed to improve the description of thiolate electrostatics and solvation, preserved a stable substrate binding pose and a persistent C256-H136 interaction, thereby tightening the catalytic site configuration. The Sγ_C256_-Nε_H136_ distance remained more stable in the CY2/HIP system than in the CYM/HIP system, further supporting the improved stability of the CY2/HIP configuration (Figure S5).

### The influence of [4Fe-4S] redox state on the active site and ligand dynamics

To assess how cluster redox state affects active site organization and reactivity, the CY2/HIP systems were analyzed with the [4Fe-4S] cluster in both oxidized and reduced states. In the oxidized [4Fe–4S]^2+^ state, coordination by four thiolate residues results in an overall charge of −2. In this system, the substrate remained stably bound across three independent 600 ns MD trajectories, as shown by the substrate RMSD profile in Figure S4. The adenosine moiety formed persistent hydrogen bonds with S60, L85, G161 and H259, whereas the negatively charged phosphosulfate group is stabilized through interactions with R242, R245, and K144 (Figure S6). The reduced CY2/HIP system, in which the [4Fe–4S]^+^ cluster carries an overall charge of −3, showed higher ligand RMSD values and increased C-terminal flexibility, as reflected by the per-residue RMSF values projected on the APSR structure (Figure 2). This increased mobility is consistent with the higher negative charge of the reduced cluster, which destabilizes the catalytic thiolate via electrostatic repulsion. Overall, the comparison of the reduced and oxidized CY2/HIP systems indicates that the oxidized [4Fe–4S]^2+^ state maintains the geometry and the electrostatic environment required for efficient nucleophilic attack. The remaining protonation configurations are reported in Figure S3-S4 for comparison.

**Figure 2.**
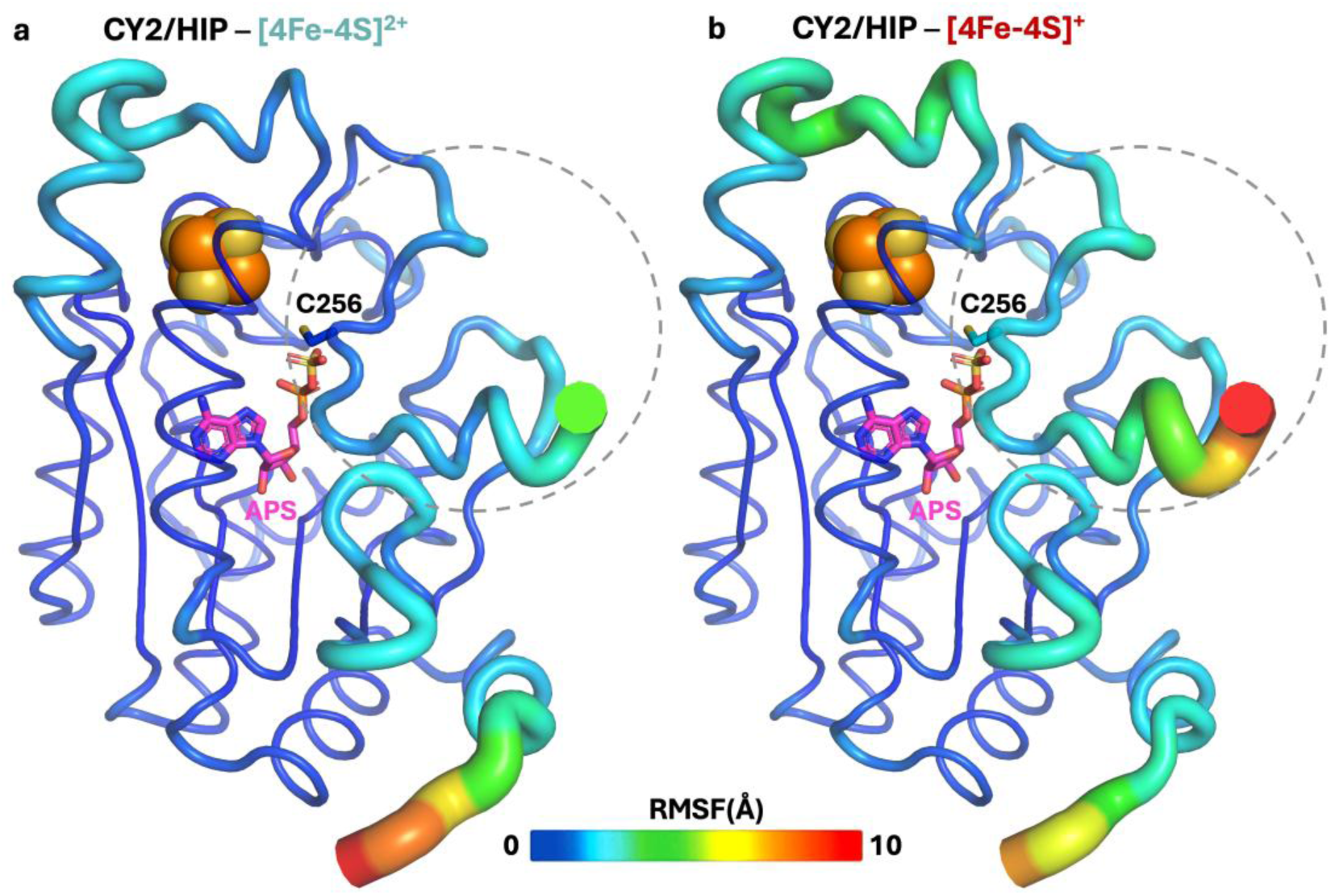
Protein Cα RMSF values are mapped on the APSR structure to visualize per-residue flexibility in CY2/HIP systems with distinct cluster redox states. RMSF values were calculated from three independent 600 ns MM MD trajectories for the oxidized [4Fe–4S]^2+^ system (a) and the reduced [4Fe– 4S]^+^ system (b). RMSF profiles showing lower C-terminus (indicated in dashed circled region) flexibility in the [4Fe–4S]^2+^ system and higher fluctuations in the [4Fe–4S]^+^ system, consistent with C256 thiolate (shown as sticks) destabilization induced by the increased negative charge of the reduced cluster.

### The influence of [4Fe-4S] redox state on catalytically relevant distances

Free energy surfaces (FESs), defined by the two catalytic distances shown in Figure 1, were generated from the MM MD trajectories. For simulations employing the validated CY2 thiolate parameters, the conformational landscapes of the CY2/HIP systems with oxidized and reduced states of the [4Fe–4S] cluster are shown in Figure 3, while those of the remaining systems are provided in Figure S7. The two redox states displayed distinct conformational behaviors. The CY2/HIP system with reduced iron-sulfur cluster sampled a broad free energy landscape, with multiple minima distributed over longer catalytic distances, particularly along the C256-H136 coordinate, indicative of higher active site flexibility and weaker interaction between C256 and H136. In contrast, the CY2/HIP system with an oxidized iron-sulfur cluster exhibited a single energy minimum. This deep minimum corresponds to a stable C256-H136 interaction and a more frequent population of sulfur–sulfur distances compatible with nucleophilic attack. The lowest free energy basins of both CY2/HIP landscapes were therefore used to guide the identification of representative starting structures for the subsequent QM/MM simulations. The selected structures were characterized by catalytic distances of *d*_1_ = 3.3 Å and *d*_2_ = 4.0 Å for the oxidized cluster, and *d*_1_ = 3.3 Å and *d*_2_ = 4.3 Å for the reduced cluster.

**Figure 3.**
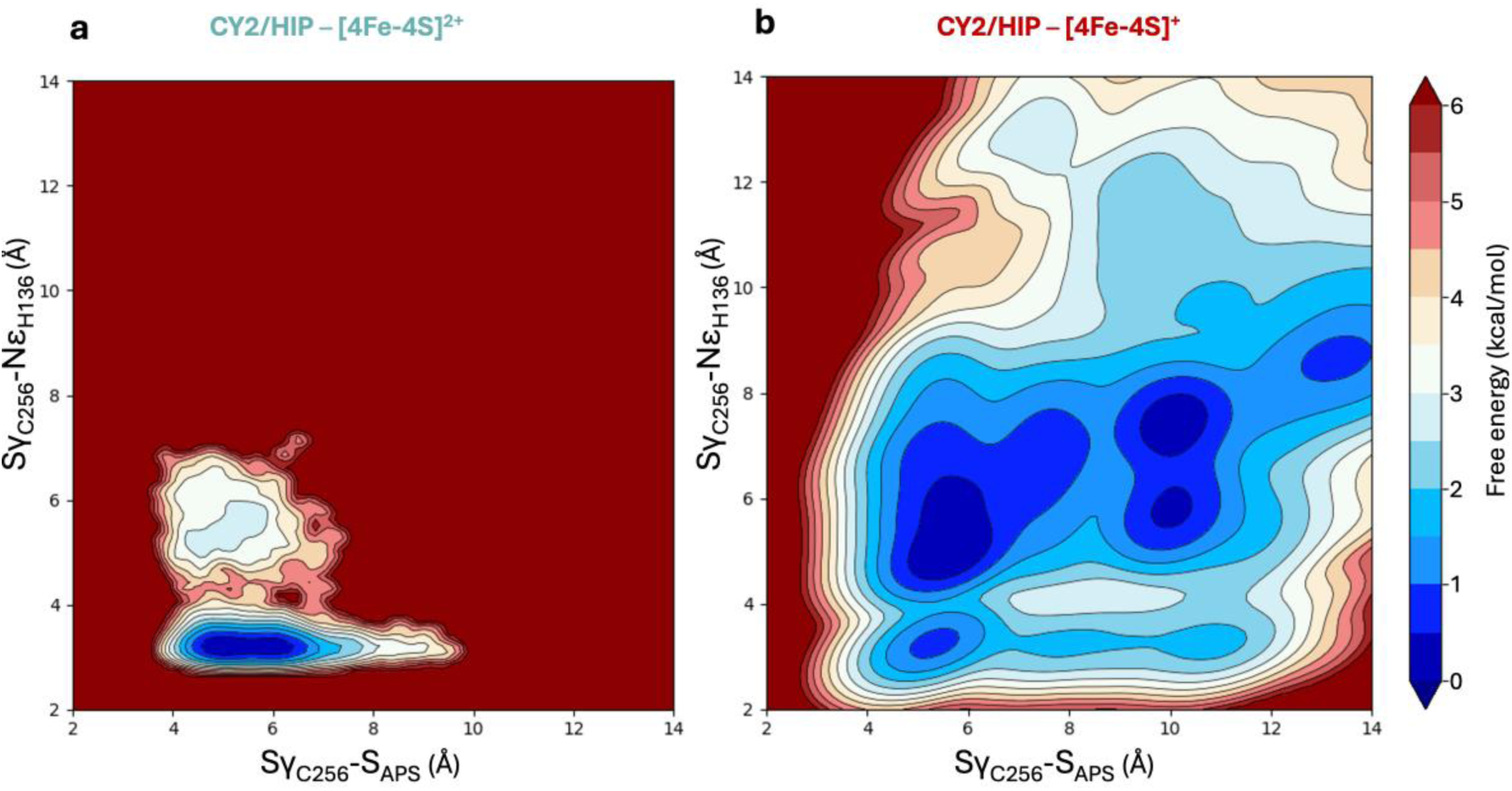
Free energy surface (FES) of the APS-APSR complex from MM MD simulations, constructed using the catalytically relevant distances *d*_1_ = Sγ_C256_-Nε_H136_ and *d*_2_ = Sγ_C256_-S_APS_. (a) The oxidized [4Fe– 4S]^2+^ system exhibits a localized low-energy basin at short catalytic distances, consistent with pre-reactive geometry. (b) The reduced [4Fe–4S]^+^ system displays multiple shallow free energy minima at longer distances, indicating diminished structural and electrostatic preorganization for catalysis.

### QM/MM simulations of the sulfite transfer

Preliminary benchmarking, reported in the Supporting Information (Sections S1–S3), indicated that PM6 provides the most accurate description of the tetrahedral geometry of the sulfur atom present in the phosphosulfate moiety (Figure S8). Additional tests identified 200 kcal mol^−1^ Å^−2^ as the optimal force constant for umbrella sampling simulations (Figure S9). Force constants above or below this value led to lack of overlap between adjacent windows along the reaction coordinate, particularly near the transition state.

Two QM-region definitions of different sizes were tested. The larger QM region was selected because the smaller model overestimated the energy barrier, while inclusion of the relevant water molecules substantially lowered the computed barrier (Figure S10). While the small QM region remains geometrically compact in the reactant state, this is no longer the case as the reaction proceeds and the nucleophilic attack begins. Specifically, the solvation pattern of the phosphate group changes significantly as the O–S bond of APS breaks and the Sγ_C256_–S_APS_ bond forms, with water molecules becoming intercalated between the phosphate and sulfite groups (Figure S11). This leads to the fragmentation of the QM region into two disconnected subregions, introducing discontinuities in the evaluation of both the QM potential energy and the QM/MM interactions, a well-known artifact in QM/MM calculations that can lead to significant errors in computed energetics.^57,58^ By contrast, when a larger QM region is employed to explicitly include the relevant water molecules, the atoms in the QM region remained closer to each other throughout the entire reaction coordinate. Therefore, the same optimized QM/MM protocol was applied to both oxidized and reduced cluster states to evaluate redox-dependent modulation of enzymatic reaction at the DFT level.

The suitability of the selected DFT level is supported by the nature of the reaction investigated here. The limitations reported for B3LYP in cysteine thio-Michael additions are associated with enolate-like intermediates in which the negative charge is delocalized across a conjugated carbonyl system.^59,60,61^ In APSR, the Cys256 thiolate attacks the sulfuryl atom of APS through nucleophilic substitution, with concomitant AMP leaving group and formation of an enzyme sulfonate intermediate. Previous studies have demonstrated the reliability of B3LYP-D3(BJ) for related nucleophilic substitution reactions at sulfur.^62,63,64,65^

The resulting PMF profiles revealed a pronounced dependence of the reaction energetics on the oxidation state of the iron-sulfur cluster. The 10 ps sampling of the oxidized [4Fe–4S]^2+^ system yielded an activation barrier of 17.7 ± 1.7 kcal mol^−1^ (Figure 4a). In contrast, reduction of the cluster penalized both the thermodynamics and kinetics of the reaction, raising the free energy of the product state and increasing the free energy barrier to 21.5 ± 2.2 kcal mol^−1^. This energetic penalty is partly a result of a less effective stabilization provided by K144 in the product state of the reduced system, compared with the more favorable hydrogen-bonding pattern observed in the oxidized system. A similar effect is also observed at the transition state. Representative structures along the reaction pathway are shown in Figure 4b and Figure 4c for the oxidized and reduced iron-sulfur cluster states, respectively.

**Figure 4.**
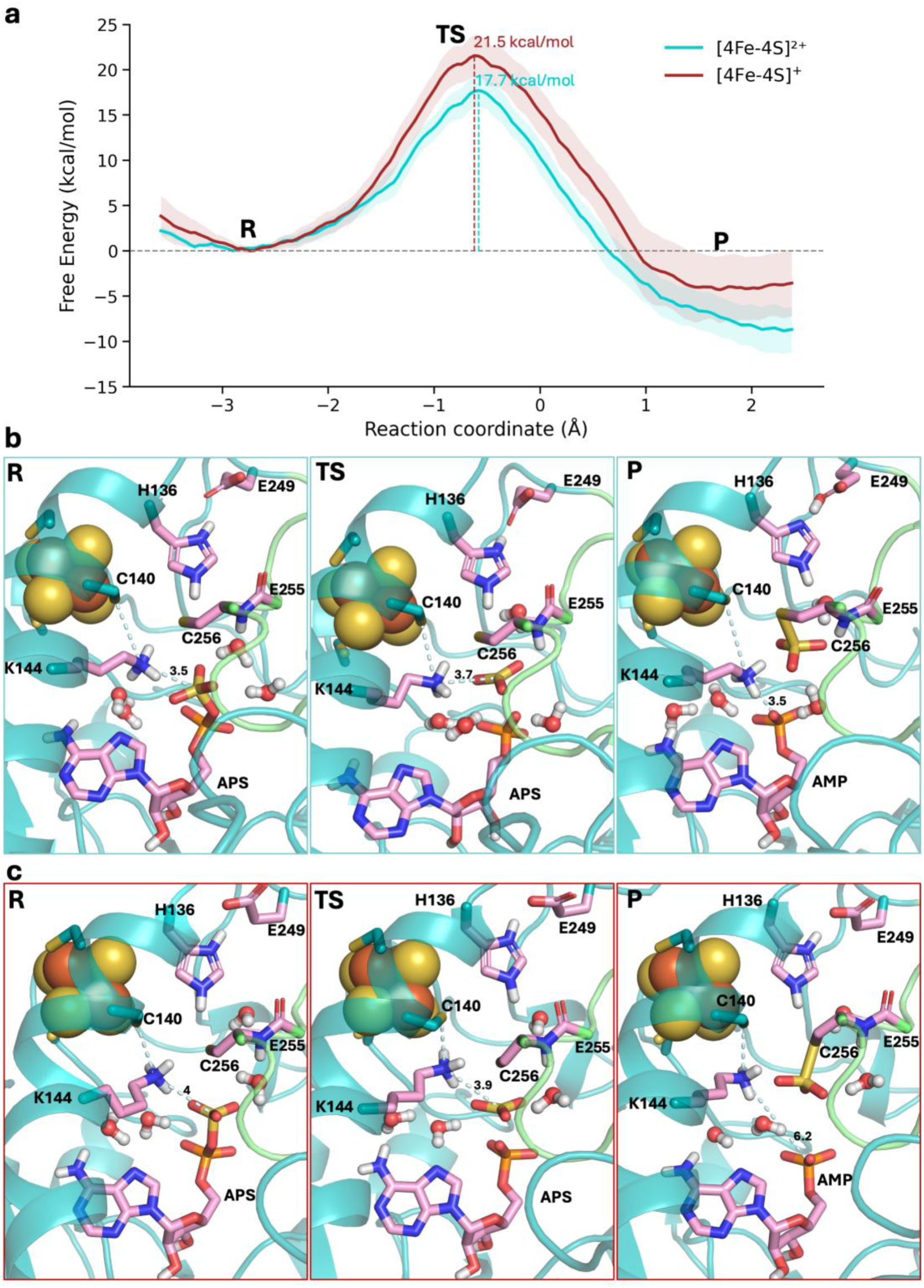
C256-mediated nucleophilic attack on APS in the oxidized [4Fe–4S]^2+^ (cyan) and reduced [4Fe-4S]^+^ (brown) cluster states. (a) Potential of mean force (PMF) from umbrella sampling QM/MM MD simulations at the DFT (B3LYP-D3(BJ)/6-31G(d)) level. Representative QM/MM snapshots along the reaction coordinate for the reactant (R), transition state (TS), and product (P), highlighting key K144 interactions presented as discontinues lines for the (b) oxidized [4Fe–4S(Cys)_4_]^2−^ and (c) reduced [4Fe–4S(Cys)_4_]^3−^ systems. The C-terminal loop is shown as a green cartoon, while QM region atoms are depicted as pink sticks. Distances are expressed in angstroms.

According to the transition state theory, the calculated barrier for APSR with the oxidized [4Fe–4S]^2^⁺ cluster agrees with the experimental activation barrier of 18.6 kcal mol⁻^1^, derived from the reported *M. tuberculosis* APSR, *k*_cat_= 9.2 ± 0.1 s^−1^ at 298.15 K,^22,23^ supporting the oxidized iron-sulfur cluster as the catalytically relevant redox state.

The lower activation barrier observed for the oxidized system arises from improved active site preorganization and hydrogen bonding, as supported by the redox-dependent changes in key active site distances shown in Figure S12.

In the oxidized system, the less negative charge supports a balanced C140-K144-APS interaction network. K144 adopts a bridging position between the cluster-coordinating C140 residue and the APS sulfate group, with interaction distances of approximately 3.5 Å to both sites. The K144 side chain stabilizes the APS sulfate group through electrostatic interactions and hydrogen bonding with the sulfate oxygen atoms, thereby attenuating repulsion between the C256 thiolate and the negatively charged substrate, promoting a catalytically competent geometry for nucleophilic attack (Figure 4b). In the reduced system, the increased negative charge of the [4Fe–4S(Cys)_4_]^3–^ site draws K144 toward the cluster and away from APS. Accordingly, K144 shifts toward C140, resulting in a K144-C140 distance of approximately 3.2 Å, while remaining around 4.0 Å from the APS sulfate group. Consequently, K144 interacts less effectively with the substrate along the reaction coordinate, including in the transition state and product, matching the increased activation barrier and reduced stabilization of the product (Figure 4c). A similar redox-dependent rearrangement is observed for H136. In the reduced system, H136 is positioned closer to the cluster-coordinating C140 residue, with an average H136-C140 distance of approximately 3.8 ± 0.15 Å, compared with 4.7 ± 0.18 Å in the oxidized system (Figure S12). This displacement places H136 farther from the catalytic C256 thiolate, potentially weakening its contribution to C256 positioning during nucleophilic attack.

This redox-dependent reorganization of the catalytic network is further supported by the transition state structure analysis. The four water molecules included in the QM region capture local solvation of the reacting group, while two of them remain associated with K144 along the reaction coordinate. In the oxidized system, these waters cooperate with K144 to form a hydrogen bonding network oriented toward the APS sulfate group, stabilizing the transition state region and facilitating sulfite transfer, further aided by the coordinating peptide bond between E255 and C256. This environment helps preserve the trigonal-planar transition state geometry and attenuates electrostatic repulsion between APS and the C256 thiolate (Figure S13). In the reduced system, the larger K144-APS separation displaces the associated waters from the reacting group, weakening sulfite stabilization along the reaction pathway.

Following S-S bond formation in the oxidized system, the K144-water network rearranges around the AMP phosphate group, where enhanced hydration and a short K144-phosphate interaction of approximately 3.5 Å contribute to product-state stabilization. By contrast, in the reduced cluster system, this distance increases to approximately 6.2 Å and the radial distribution function (RDF) analysis reveals a less ordered hydration shell, consistent with weaker product stabilization (Figure S11).

The RDF of water molecules around the phosphorus and sulfur atoms reveals distinct solvation between the reactant state (APS) and the product state (AMP and thiosulfate). In the product state, water molecules are positioned around the AMP phosphate group via stronger interactions, in agreement with enhanced product stabilization in the oxidized system, whereas the reduced system shows a weaker recruitment of water molecules from the bulk. In the reactant structures, these differences are less pronounced; however, the APS sulfite group displays lower solvation in the reduced cluster than in the oxidized one. This behavior may be related to the stronger K144-sulfite hydrogen bond observed in the oxidized cluster, which may polarize the sulfite group and enhance its interaction with surrounding water molecules. This can also be seen through the analysis of the dipole moment of the sulfite with the two oxidation states of the iron-sulfur cluster (Figure S14). This electrostatic effect may therefore account for the greater sulfite solvation observed in the oxidized system relative to the reduced one.

The local electric field calculations were performed using the TUPÃ program (see Section S4 in the SI for details).^66^ Oxidized system generates a substantially stronger electric field at the substrate oxygen involved in the cleavage of the phosphate-sulfate bond (15–20 MV/cm larger in R and TS; ∼177 MV/cm larger in P) compared to the reduced state system, indicating enhanced electrostatic preorganization that stabilizes the developing negative charge along the reaction coordinate (Figure S15). Per-residue electric field decomposition identifies K144 as the primary contributor to this stabilization (Figure S16), suggesting that its electrostatic interaction with the substrate is redox-tuned by the cluster. This stronger field in the oxidized state lowers the activation barrier and provides greater thermodynamic driving force, predicting faster reaction kinetics in the oxidized state consistent with an electric field-driven catalytic mechanism.

Overall, these findings indicate that the oxidized cluster helps maintain the electrostatic organization required for efficient C256-mediated nucleophilic attack, whereas cluster reduction disrupts this organization, weakens transition state and product stabilization, and ultimately raises the activation barrier. Taken together, our QM/MM simulations support two complementary catalytic roles for K144: (i) it promotes a reactive orientation of the APS sulfate group for nucleophilic attack by C256, (ii) it coordinates active site waters that contribute to stabilization of both transition state and product. Consistent with this mechanistic role, mutation of K144 to alanine results in a pronounced loss of catalytic activity, confirming its direct contribution to catalysis.^67^ In line with this interpretation, Bhave et al. combined DFT calculations with X-ray absorption spectroscopy to show that K144 mediates electrostatic coupling between the iron-sulfur cluster and APS, thereby promoting phosphosulfate stabilization.^39^ A similar role has been reported before for the conserved lysine in sulfotransferases, which stabilizes the PAPS transition state through electrostatic engagement of the transferring SO_3_^2⁻^ group.^68^

## CONCLUSIONS

APSR catalyzes the essential first step of assimilatory sulfate reduction in *Pseudomonas aeruginosa* by transferring the sulfite group from adenosine 5’-phosphosulfate (APS) to a catalytic cysteine, generating AMP and an enzyme-bound S-sulfocysteine intermediate.

To elucidate this mechanism from an incomplete crystal structure, we modeled the missing C-terminal tail using AlphaFold, which successfully reproduced the catalytic cysteine geometry observed in the homologous PAPS reductase structure.^14^ Extensive MM MD simulations identified catalytically relevant protonation states for the catalytic H136/C256 pair and highlighted the importance of proper parameterization of the C256 thiolate, required to accurately capture solvent contacts and preserve a catalytically competent active site environment. The MD analyses further showed that the oxidized [4Fe–4S]^2+^ cluster maintains stable substrate binding, limits C-terminus flexibility, and favors compact catalytic geometries. By contrast, the reduced [4Fe–4S]^+^ state increases substrate mobility and active site flexibility, with increased fluctuations of the catalytic residues. Together, these results indicate that the oxidized cluster provides the electrostatic environment required for catalysis, whereas the reduced state generates an overly negative electrostatic potential in the active site that perturbs the catalytic thiolate, ligand solvation and product stabilization, consistent with experimental evidence supporting the oxidized cluster as the catalytically competent redox state.^18^

QM/MM MD umbrella sampling simulations provided the free energy profile for C256-mediated nucleophilic attack. The DFT QM/MM MD simulations at the B3LYP-D3(BJ)/6-31G(d) level capture the key geometric features of the reaction coordinate and provide an activation free energy of 17.7 ± 1.7 kcal mol^−1^, which is in good agreement with the barrier derived from the *k*_cat_ measured for the *M. tuberculosis* ortholog.^22,23^ We demonstrate that the oxidation state of the [4Fe–4S] cluster modulates the electrostatic organization of the catalytic site, with the oxidized state maintaining K144 in a catalytically productive position and supporting the geometry and electrostatics required for nucleophilic attack. The conserved K144 residue plays an important mechanistic role by stabilizing the sulfate group of APS through the transition state into the phosphate group of the AMP product. Active site water molecules further contribute by solvating the negatively charged phosphosulfate group of APS in reactant state, reducing electrostatic repulsion with the C256 thiolate and thereby the activation barrier, while also stabilizing the AMP product formed.

Our mechanistic study establishes the molecular basis for catalysis in *P. aeruginosa* APSR, identifying the roles of the oxidized [4Fe–4S] cluster, K144, C256 and active site waters in stabilizing the transition state. Specifically, our simulations reveal the closed conformation of the C-terminal loop and define the arrangement of catalytic residues in the Michaelis complex, providing a structurally and electrostatically detailed picture of the active site that has not previously been available for this ortholog. This mechanistic rationale underscores the distinct electrostatic environment of the active site, shaped by the oxidized cluster and the charge-stabilizing role of K144, defining hotspots for substrate interaction.

This work provides a validated structural framework for drug discovery, given that existing APSR inhibitors are largely nucleoside mimetics^69,70^ or weak hits developed against the *M. tuberculosis* ortholog.^71,72^ By providing a high-resolution mechanistic description of this clinically relevant antimicrobial target, our work supports the rational design of covalent and non-covalent inhibitors with improved potency and selectivity. These findings suggest that the redox-dependent organization of the active site may influence inhibitor recognition and covalent engagement of C256. In this context, interactions with K144 represent a potentially exploitable feature for the structure-based design of APSR inhibitors. More broadly, by defining how the APSR active site and its [4Fe–4S] cluster respond to distinct redox states, this work highlights the importance of redox control within the assimilatory sulfur-reduction pathway. Targeting this redox-sensitive catalytic machinery may therefore provide an additional strategy for disrupting sulfur metabolism and advancing antimicrobial development against antibiotic-resistant infections.

## ASSOCIATED CONTENT

### Table Of Contents

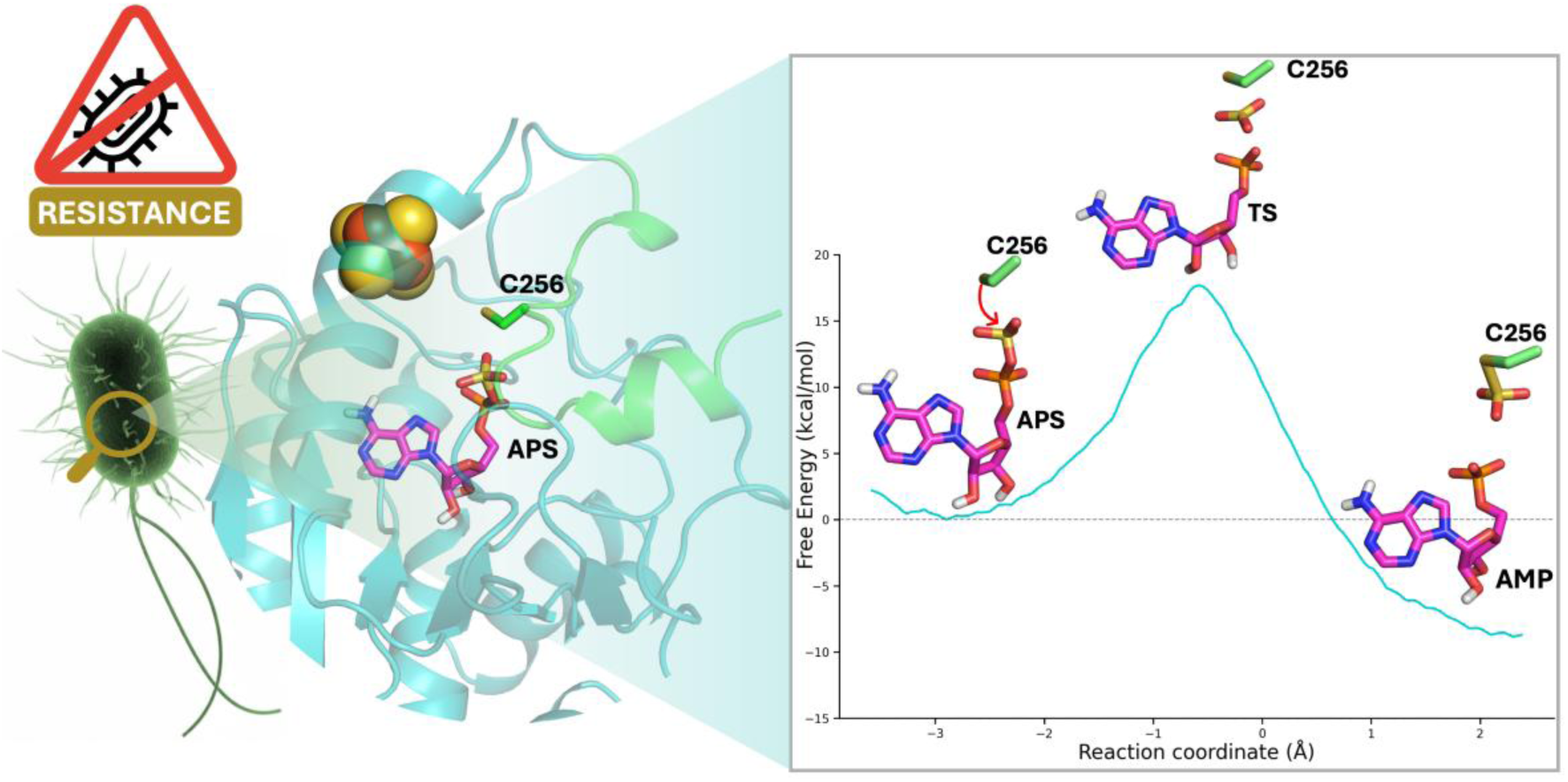

### Supporting Information

The Supporting Information is available free of charge at https://pubs.acs.org/.

Additional data analysis is provided in supplementary figures S1-S16 and sections S1-S4.

## DATA AVAILABILITY

The AMBER topologies, coordinates, input files, representative MM and QM/MM MD trajectories, and raw PMF data are deposited on Zenodo: 10.5281/zenodo.22098422

## ACKNOWLEDGMENT

We gratefully acknowledge support from the EPSRC through the grant *Predictive multiscale free energy simulations of hybrid transition metal catalysis* (FEHy-bCat; EP/W013738/1). We also thank the European Research Council for funding under the European Union’s Horizon 2020 research and innovation program (PREDACTED Advanced Grant Agreement No. 101021207). UK Catalysis Hub is kindly thanked for resources and support provided via our membership of the UK Catalysis Hub Consortium and funded by EPSRC grant: UKRI945. This work used the computational and data storage facilities of the Advanced Computing Research Centre at the University of Bristol. Access to the Isambard 3 Tier-2 HPC Facility was also provided; Isambard 3 is hosted by the University of Bristol, operated by the GW4 Alliance, and funded by UK Research and Innovation and the Engineering and Physical Sciences Research Council under grant EP/X039137/1. We acknowledge ISCRA for awarding this project access to the LEONARDO supercomputer at CINECA, Italy, through the project HP10CSXV7W. This work has been also funded by the European Union─NextGenerationEU, Mission 4, Component 2, under the Italian Ministry of University and Research (MUR) National Innovation Ecosystem grant ECS00000041─VITALITY─CUP H33C22000430006.

## REFERENCES

(1) Ho, C. S.; Wong, C. T. H.; Aung, T. T.; Lakshminarayanan, R.; Mehta, J. S.; Rauz, S.; McNally, A.; Kintses, B.; Peacock, S. J.; de la Fuente-Nunez, C.; Hancock, R. E. W.; Ting, D. S. J. Antimicrobial Resistance: A Concise Update. Lancet Microbe 2025, 6 (1), 100947. 10.1016/j.lanmic.2024.07.010

(2) Boucher, H. W.; Talbot, G. H.; Bradley, J. S.; Edwards, J. E.; Gilbert, D.; Rice, L. B.; Scheld, M.; Spellberg, B.; Bartlett, J. Bad Bugs, No Drugs: No ESKAPE! An Update from the Infectious Diseases Society of America. Clinical Infectious Diseases 2009, 48 (1), 1–12. 10.1086/595011

(3) Malhotra, S.; Hayes, D.; Wozniak, D. J. Cystic Fibrosis and Pseudomonas Aeruginosa: The Host-Microbe Interface. Clin. Microbiol. Rev. 2019, 32 (3). 10.1128/CMR.00138-18

(4) Tralau, T.; Vuilleumier, S.; Thibault, C.; Campbell, B. J.; Hart, C. A.; Kertesz, M. A. Transcriptomic Analysis of the Sulfate Starvation Response of *Pseudomonas Aeruginosa*. J. Bacteriol. 2007, 189 (19), 6743–6750. 10.1128/JB.00889-07

(5) Tao, Y.; Zheng, D.; Zou, W.; Guo, T.; Liao, G.; Zhou, W. Targeting the Cysteine Biosynthesis Pathway in Microorganisms: Mechanism, Structure, and Drug Discovery. Eur. J. Med. Chem. 2024, 271, 116461. 10.1016/j.ejmech.2024.116461

(6) Spigelmyer, S. M.; Dos Santos, P. C. Intricacies in Iron–Sulfur Cluster Function and Biogenesis: Functional Versatility, Sulfur Sources, and Enzyme Specificity. RSC Chem. Biol. 2026, 7 (5), 763–782. 10.1039/D5CB00330J

(7) Carroll, K. S.; Gao, H.; Chen, H.; Stout, C. D.; Leary, J. A.; Bertozzi, C. R. A Conserved Mechanism for Sulfonucleotide Reduction. PLoS Biol. 2005, 3 (8), e250. 10.1371/journal.pbio.0030250

(8) Feliciano, P. R.; Carroll, K. S.; Drennan, C. L. Crystal Structure of the [4Fe–4S] Cluster-Containing Adenosine-5′-Phosphosulfate Reductase from *Mycobacterium Tuberculosis*. ACS Omega 2021, 6 (21), 13756–13765. 10.1021/acsomega.1c01043

(9) Chartron, J.; Carroll, K. S.; Shiau, C.; Gao, H.; Leary, J. A.; Bertozzi, C. R.; Stout, C. D. Substrate Recognition, Protein Dynamics, and Iron-Sulfur Cluster in Pseudomonas Aeruginosa Adenosine 5′-Phosphosulfate Reductase. J. Mol. Biol. 2006, 364 (2), 152–169. 10.1016/j.jmb.2006.08.080

(10) Chartron, J.; Shiau, C.; Stout, C. D.; Carroll, K. S. 3‘-Phosphoadenosine-5‘-Phosphosulfate Reductase in Complex with Thioredoxin: A Structural Snapshot in the Catalytic Cycle,. Biochemistry 2007, 46 (13), 3942–3951. 10.1021/bi700130e

(11) Palde, P. B.; Carroll, K. S. A Universal Entropy-Driven Mechanism for Thioredoxin–Target Recognition. Proceedings of the National Academy of Sciences 2015, 112 (26), 7960–7965. 10.1073/pnas.1504376112

(12) Hong, J. A.; Carroll, K. S. Deciphering the Role of Histidine 252 in Mycobacterial Adenosine 5′-Phosphosulfate (APS) Reductase Catalysis. Journal of Biological Chemistry 2011, 286 (32), 28567–28573. 10.1074/jbc.M111.238998

(13) Chartron, J., C. K. S., S. C., G. H., L. J. A., B. C. R., S. C. D. Crystal Structure of Assimilatory Adenosine 5’-Phosphosulfate Reductase with Bound APS. Journal of Molecular Biology 2006. 10.2210/pdb2goy/pdb

(14) Yu, Z.; Lemongello, D.; Segel, I. H.; Fisher, A. J. Crystal Structure of *Saccharomyces Cerevisiae* 3′-Phosphoadenosine-5′-Phosphosulfate Reductase Complexed with Adenosine 3′,5′-Bisphosphate. Biochemistry 2008, 47 (48), 12777–12786. 10.1021/bi801118f

(15) Kim, S.-K.; Rahman, A.; Mason, J. T.; Hirasawa, M.; Conover, R. C.; Johnson, M. K.; Miginiac-Maslow, M.; Keryer, E.; Knaff, D. B.; Leustek, T. The Interaction of 5′-Adenylylsulfate Reductase from Pseudomonas Aeruginosa with Its Substrates. Biochimica et Biophysica Acta (BBA) - Bioenergetics 2005, 1710 (2–3), 103–112. 10.1016/j.bbabio.2005.09.004

(16) Carroll, K. S.; Gao, H.; Chen, H.; Leary, J. A.; Bertozzi, C. R. Investigation of the Iron−Sulfur Cluster in *Mycobacterium Tuberculosis* APS Reductase: Implications for Substrate Binding and Catalysis. Biochemistry 2005, 44 (44), 14647–14657. 10.1021/bi051344a

(17) Bhave, D. P.; Hong, J. A.; Keller, R. L.; Krebs, C.; Carroll, K. S. Iron–Sulfur Cluster Engineering Provides Insight into the Evolution of Substrate Specificity among Sulfonucleotide Reductases. ACS Chem. Biol. 2012, 7 (2), 306–315. 10.1021/cb200261n

(18) Gao, H.; Leary, J.; Carroll, K. S.; Bertozzi, C. R.; Chen, H. Noncovalent Complexes of APS Reductase from *M. Tuberculosis* : Delineating a Mechanistic Model Using ESI-FTICR MS. J. Am. Soc. Mass Spectrom. 2007, 18 (2), 167–178. 10.1016/j.jasms.2006.08.010

(19) Hanževački, M.; Croft, A. K.; Jäger, C. M. Activation of Glycyl Radical Enzymes─Multiscale Modeling Insights into Catalysis and Radical Control in a Pyruvate Formate-Lyase-Activating Enzyme. J. Chem. Inf. Model. 2022, 62 (14), 3401–3414. 10.1021/acs.jcim.2c00362

(20) Nicolet, Y. Structure–Function Relationships of Radical SAM Enzymes. Nat. Catal. 2020, 3 (4), 337–350. 10.1038/s41929-020-0448-7

(21) Jia, T.; Peng, Y.; Niu, L.; Qi, Z.; Xi, J. Simultaneous Sulfide Oxidation and Sulfate Reduction for Intracellular Redox Homeostasis under Highly Acidic Conditions. Nat. Commun. 2026, 17 (1), 1797. 10.1038/s41467-026-68508-y

(22) Sun, M.; Leyh, T. S. Channeling in Sulfate Activating Complexes. Biochemistry 2006, 45 (38), 11304–11311. 10.1021/bi060421e

(23) Paritala, H.; Carroll, K. S. A Continuous Spectrophotometric Assay for Adenosine 5′-Phosphosulfate Reductase Activity with Sulfite-Selective Probes. Anal. Biochem. 2013, 440 (1), 32–39. 10.1016/j.ab.2013.05.007

(24) Madhavi Sastry, G.; Adzhigirey, M.; Day, T.; Annabhimoju, R.; Sherman, W. Protein and Ligand Preparation: Parameters, Protocols, and Influence on Virtual Screening Enrichments. J. Comput. Aided. Mol. Des. 2013, 27 (3), 221–234. 10.1007/s10822-013-9644-8

(25) Jacobson, M. P.; Pincus, D. L.; Rapp, C. S.; Day, T. J. F.; Honig, B.; Shaw, D. E.; Friesner, R. A. A Hierarchical Approach to All-atom Protein Loop Prediction. Proteins: Structure, Function, and Bioinformatics 2004, 55 (2), 351–367. 10.1002/prot.10613

(26) Jumper, J.; Evans, R.; Pritzel, A.; Green, T.; Figurnov, M.; Ronneberger, O.; Tunyasuvunakool, K.; Bates, R.; Žídek, A.; Potapenko, A.; Bridgland, A.; Meyer, C.; Kohl, S. A. A.; Ballard, A. J.; Cowie, A.; Romera-Paredes, B.; Nikolov, S.; Jain, R.; Adler, J.; Back, T.; Petersen, S.; Reiman, D.; Clancy, E.; Zielinski, M.; Steinegger, M.; Pacholska, M.; Berghammer, T.; Bodenstein, S.; Silver, D.; Vinyals, O.; Senior, A. W.; Kavukcuoglu, K.; Kohli, P.; Hassabis, D. Highly Accurate Protein Structure Prediction with AlphaFold. Nature 2021, 596 (7873), 583–589. 10.1038/s41586-021-03819-2

(27) Lu, C.; Wu, C.; Ghoreishi, D.; Chen, W.; Wang, L.; Damm, W.; Ross, G. A.; Dahlgren, M. K.; Russell, E.; Von Bargen, C. D.; Abel, R.; Friesner, R. A.; Harder, E. D. OPLS4: Improving Force Field Accuracy on Challenging Regimes of Chemical Space. J. Chem. Theory Comput. 2021, 17 (7), 4291–4300. 10.1021/acs.jctc.1c00302

(28) Gordon, J. C.; Myers, J. B.; Folta, T.; Shoja, V.; Heath, L. S.; Onufriev, A. H++: A Server for Estimating PKas and Adding Missing Hydrogens to Macromolecules. Nucleic Acids Res. 2005, 33 (Web Server), W368–W371. 10.1093/nar/gki464

(29) Maier, J. A.; Martinez, C.; Kasavajhala, K.; Wickstrom, L.; Hauser, K. E.; Simmerling, C. Ff14SB: Improving the Accuracy of Protein Side Chain and Backbone Parameters from Ff99SB. J. Chem. Theory Comput. 2015, 11 (8), 3696–3713. 10.1021/acs.jctc.5b00255

(30) Frisch, M. J. ;T. G. W.; S. H. B. ; S. G. E. ; R. M. A. ; C. J. R. ; S. G. ; B. V. ; P. G. A. ; N. H.;, et al. G. Gaussian 16. Revision C.01; Gaussian, Inc.: Wallingford, CT, 2016.

(31) Wang, J.; Wolf, R. M.; Caldwell, J. W.; Kollman, P. A.; Case, D. A. Development and Testing of a General Amber Force Field. J. Comput. Chem. 2004, 25 (9), 1157–1174. 10.1002/jcc.20035

(32) Bayly, C. I.; Cieplak, P.; Cornell, W.; Kollman, P. A. A Well-Behaved Electrostatic Potential Based Method Using Charge Restraints for Deriving Atomic Charges: The RESP Model. J. Phys. Chem. 1993, 97 (40), 10269–10280. 10.1021/j100142a004

(33) Li, P.; Song, L. F.; Merz, K. M. Systematic Parameterization of Monovalent Ions Employing the Nonbonded Model. J. Chem. Theory Comput. 2015, 11 (4), 1645–1657. 10.1021/ct500918t

(34) Jorgensen, W. L.; Chandrasekhar, J.; Madura, J. D.; Impey, R. W.; Klein, M. L. Comparison of Simple Potential Functions for Simulating Liquid Water. J. Chem. Phys. 1983, 79 (2), 926–935. 10.1063/1.445869

(35) Case, D. A.; Aktulga, H. M.; Belfon, K.; Cerutti, D. S.; Cisneros, G. A.; Cruzeiro, V. W. D.; Forouzesh, N.; Giese, T. J.; Götz, A. W.; Gohlke, H.; Izadi, S.; Kasavajhala, K.; Kaymak, M. C.; King, E.; Kurtzman, T.; Lee, T.-S.; Li, P.; Liu, J.; Luchko, T.; Luo, R.; Manathunga, M.; Machado, M. R.; Nguyen, H. M.; O’Hearn, K. A.; Onufriev, A. V.; Pan, F.; Pantano, S.; Qi, R.; Rahnamoun, A.; Risheh, A.; Schott-Verdugo, S.; Shajan, A.; Swails, J.; Wang, J.; Wei, H.; Wu, X.; Wu, Y.; Zhang, S.; Zhao, S.; Zhu, Q.; Cheatham, T. E.; Roe, D. R.; Roitberg, A.; Simmerling, C.; York, D. M.; Nagan, M. C.; Merz, K. M. AmberTools. J. Chem. Inf. Model. 2023, 63 (20), 6183–6191. 10.1021/acs.jcim.3c01153

(36) Li, P.; Merz, K. M. MCPB.Py: A Python Based Metal Center Parameter Builder. J. Chem. Inf. Model. 2016, 56 (4), 599–604. 10.1021/acs.jcim.5b00674

(37) Seminario, J. M. Calculation of Intramolecular Force Fields from Second-Derivative Tensors. Int. J. Quantum Chem. 1996, 60 (7), 1271–1277. 10.1002/(SICI)1097-461X(1996)60:7<1271::AID-QUA8>3.0.CO;2-W

(38) Landry, L.; Li, P. Parameterization of a Fluctuating Charge Model for Complexes Containing 3d Transition Metals. J. Phys. Chem. B 2024, 128 (42), 10329–10338. 10.1021/acs.jpcb.4c03219

(39) Bhave, D. P.; Han, W.-G.; Pazicni, S.; Penner-Hahn, J. E.; Carroll, K. S.; Noodleman, L. Geometric and Electrostatic Study of the [4Fe-4S] Cluster of Adenosine-5′-Phosphosulfate Reductase from Broken Symmetry Density Functional Calculations and Extended X-Ray Absorption Fine Structure Spectroscopy. Inorg. Chem. 2011, 50 (14), 6610–6625. 10.1021/ic200446c

(40) Pedron, F. N.; Messias, A.; Zeida, A.; Roitberg, A. E.; Estrin, D. A. Novel Lennard-Jones Parameters for Cysteine and Selenocysteine in the AMBER Force Field. J. Chem. Inf. Model. 2023, 63 (2), 595–604. 10.1021/acs.jcim.2c01104

(41) Feller, S. E.; Zhang, Y.; Pastor, R. W.; Brooks, B. R. Constant Pressure Molecular Dynamics Simulation: The Langevin Piston Method. J. Chem. Phys. 1995, 103 (11), 4613–4621. 10.1063/1.470648

(42) Berendsen, H. J. C.; Postma, J. P. M.; van Gunsteren, W. F.; DiNola, A.; Haak, J. R. Molecular Dynamics with Coupling to an External Bath. J. Chem. Phys. 1984, 81 (8), 3684–3690. 10.1063/1.448118

(43) Amber 2025, U. of C. S. F. Amber 2025, University of California, San Francisco.

(44) Salomon-Ferrer, R.; Götz, A. W.; Poole, D.; Le Grand, S.; Walker, R. C. Routine Microsecond Molecular Dynamics Simulations with AMBER on GPUs. 2. Explicit Solvent Particle Mesh Ewald. J. Chem. Theory Comput. 2013, 9 (9), 3878–3888. 10.1021/ct400314y

(45) Ryckaert, J.-P.; Ciccotti, G.; Berendsen, H. J. C. Numerical Integration of the Cartesian Equations of Motion of a System with Constraints: Molecular Dynamics of n-Alkanes. J. Comput. Phys. 1977, 23 (3), 327–341. 10.1016/0021-9991(77)90098-5

(46) Darden, T.; York, D.; Pedersen, L. An N·log(N) Method for Ewald Sums in Large Systems. J. Chem. Phys. 1993, 98 (12), 10089–10092. 10.1063/1.464397

(47) https://github.com/sulfierry/free_energy_landscape.

(48) Nam, K.; Gao, J.; York, D. M. An Efficient Linear-Scaling Ewald Method for Long-Range Electrostatic Interactions in Combined QM/MM Calculations. J. Chem. Theory Comput. 2005, 1 (1), 2–13. 10.1021/ct049941i

(49) Torrie, G. M.; Valleau, J. P. Nonphysical Sampling Distributions in Monte Carlo Free-Energy Estimation: Umbrella Sampling. J. Comput. Phys. 1977, 23 (2), 187–199. 10.1016/0021-9991(77)90121-8

(50) Stewart, J. J. P. Optimization of Parameters for Semiempirical Methods V: Modification of NDDO Approximations and Application to 70 Elements. J. Mol. Model. 2007, 13 (12), 1173–1213. 10.1007/s00894-007-0233-4

(51) Grimme, S.; Antony, J.; Ehrlich, S.; Krieg, H. A Consistent and Accurate Ab Initio Parametrization of Density Functional Dispersion (DFT-D) for the 94 Elements H−Pu. J. Chem. Phys. 2010, 132 (15). 10.1063/1.3382344

(52) Lonsdale, R.; Harvey, J. N.; Mulholland, A. J. Effects of Dispersion in Density Functional Based Quantum Mechanical/Molecular Mechanical Calculations on Cytochrome P450 Catalyzed Reactions. J. Chem. Theory Comput. 2012, 8 (11), 4637–4645. 10.1021/ct300329h

(53) Manathunga, M.; Shajan, A.; Smith, J.; Miao, Y.; He, X.; Ayers, K.; Brothers, E.; Goetz, A. W.; Merz, K. M. QUICK-24.03; University of California San Diego, CA and Michigan State University: East Lansing, MI, 2024.

(54) Manathunga, M.; Aktulga, H. M.; Götz, A. W.; Merz, K. M. Quantum Mechanics/Molecular Mechanics Simulations on NVIDIA and AMD Graphics Processing Units. J. Chem. Inf. Model. 2023, 63, 711−717.

(55) Kumar, S.; Rosenberg, J. M.; Bouzida, D.; Swendsen, R. H.; Kollman, P. A. THE Weighted Histogram Analysis Method for Free-energy Calculations on Biomolecules. I. The Method. J. Comput. Chem. 1992, 13 (8), 1011–1021. 10.1002/jcc.540130812

(56) Grossfield, A. WHAM: The Weighted Histogram Analysis Method. Version 2.0.11; University of Rochester: Rochester, NY, 2019. http://Membrane.Urmc.Rochester.Edu/Wordpress/?Page_id=126.

(57) Senn, H. M.; Thiel, W. QM/MM Methods for Biomolecular Systems. Angew. Chem. Int. Ed. 2009, 48 (7), 1198–1229. 10.1002/anie.200802019

(58) Lonsdale, R.; Harvey, J. N.; Mulholland, A. J. A Practical Guide to Modelling Enzyme-Catalysed Reactions. Chem. Soc. Rev. 2012, 41 (8), 3025. 10.1039/c2cs15297e

(59) Sharma, J.; Champagne, P. A. Benchmark of Density Functional Theory Methods for the Study of Organic Polysulfides. J. Comput. Chem. 2022, 43 (32), 2131–2138. 10.1002/jcc.27007

(60) Smith, J. M.; Jami Alahmadi, Y.; Rowley, C. N. Range-Separated DFT Functionals Are Necessary to Model Thio-Michael Additions. J. Chem. Theory Comput. 2013, 9 (11), 4860–4865. 10.1021/ct400773k

(61) Awoonor-Williams, E.; Isley, W. C.; Dale, S. G.; Johnson, E. R.; Yu, H.; Becke, A. D.; Roux, B.; Rowley, C. N. Quantum Chemical Methods for Modeling Covalent Modification of Biological Thiols. J. Comput. Chem. 2020, 41 (5), 427–438. 10.1002/jcc.26064

(62) Ramos-Guzmán, C. A.; Ruiz-Pernía, J. J.; Tuñón, I. Multiscale Simulations of SARS-CoV-2 3CL Protease Inhibition with Aldehyde Derivatives. Role of Protein and Inhibitor Conformational Changes in the Reaction Mechanism. ACS Catal. 2021, 11 (7), 4157–4168. 10.1021/acscatal.0c05522

(63) Schillings, J.; Ramos-Guzmán, C. A.; Ruiz-Pernía, J. J.; Tuñón, I. Pomotrelvir and Nirmatrelvir Binding and Reactivity with SARS-CoV-2 Main Protease: Implications for Resistance Mechanisms from Computations. Angew. Chem. Int. Ed. 2024, 63 (40). 10.1002/anie.202409527

(64) Ramos-Guzmán, C. A.; Ruiz-Pernía, J. J.; Zinovjev, K.; Tuñón, I. Unveiling the Mechanistic Singularities of Caspases: A Computational Analysis of the Reaction Mechanism in Human Caspase-1. ACS Catal. 2023, 13 (7), 4348–4361. 10.1021/acscatal.3c00037

(65) Ramos-Guzmán, C. A.; Ruiz-Pernía, J. J.; Tuñón, I. Inhibition Mechanism of SARS-CoV-2 Main Protease with Ketone-Based Inhibitors Unveiled by Multiscale Simulations: Insights for Improved Designs**. Angew. Chem. Int. Ed. 2021, 60 (49), 25933–25941. 10.1002/anie.202110027

(66) Polêto, M. D.; Lemkul, J. A. TUPÃ: Electric Field Analyses for Molecular Simulations. J. Comput. Chem. 2022, 43 (16), 1113–1119. 10.1002/jcc.26873

(67) Bhave, D. P.; Hong, J. A.; Lee, M.; Jiang, W.; Krebs, C.; Carroll, K. S. Spectroscopic Studies on the [4Fe-4S] Cluster in Adenosine 5′-Phosphosulfate Reductase from Mycobacterium Tuberculosis. Journal of Biological Chemistry 2011, 286 (2), 1216–1226. 10.1074/jbc.M110.193722

(68) Kakuta, Y.; Petrotchenko, E. V.; Pedersen, L. C.; Negishi, M. The Sulfuryl Transfer Mechanism. Journal of Biological Chemistry 1998, 273 (42), 27325–27330. 10.1074/jbc.273.42.27325

(69) Hong, J. A.; Bhave, D. P.; Carroll, K. S. Identification of Critical Ligand Binding Determinants in *Mycobacterium Tuberculosis* Adenosine-5′-Phosphosulfate Reductase. J. Med. Chem. 2009, 52 (17), 5485–5495. 10.1021/jm900728u

(70) Paritala, H.; Suzuki, Y.; Carroll, K. S. Design, Synthesis and Evaluation of Fe-S Targeted Adenosine 5′-Phosphosulfate Reductase Inhibitors. Nucleosides Nucleotides Nucleic Acids 2015, 34 (3), 199–220. 10.1080/15257770.2014.978012

(71) Palde, P. B.; Bhaskar, A.; Pedró Rosa, L. E.; Madoux, F.; Chase, P.; Gupta, V.; Spicer, T.; Scampavia, L.; Singh, A.; Carroll, K. S. First-in-Class Inhibitors of Sulfur Metabolism with Bactericidal Activity against Non-Replicating *M. Tuberculosis*. ACS Chem. Biol. 2016, 11 (1), 172–184. 10.1021/acschembio.5b00517

72. A. Cosconati, S.; Hong, J. A.; Novellino, E.; Carroll, K. S.; Goodsell, D. S.; Olson, A. J. Structure-Based Virtual Screening and Biological Evaluation of *Mycobacterium Tuberculosis* Adenosine 5′-Phosphosulfate Reductase Inhibitors. J. Med. Chem. 2008, 51 (21), 6627–6630. 10.1021/jm800571m

