## Supporting Information for "Redox Control of S-sulfocysteine Formation in Adenosine Phosphosulfate Reductase"

### Table of Contents

*Figure S1. Sequence alignment and structural comparison of APSR and PAPSR.* **3**

*Figure S2. Definition of QM regions used in QM/MM simulations.* **4**

*Figure S3. Comparative protein Cα RMSF analysis of APSR across different protonation C256/H136 and [4Fe–4S] cluster oxidation states* **5**

*Figure S4. Comparative APS RMSD analysis of APSR in distinct protonation and [4Fe–4S] cluster oxidation states.* **6**

*Figure S5. Thiolate parameters affect active site stability and hydration* **7**

*Figure S6. APSR-substrate hydrogen bond interaction frequency in the CY2/HIP oxidized* **7**

*Figure S7. Free energy landscapes of APSR C256/H136 protonation states.* **8**

*Section S1. Benchmarking of semiempirical QM/MM methods* **9**

*Figure S8. Semiempirical description of APS phosphosulfate geometry.* **9**

*Section S2. Optimization of the umbrella sampling protocol* **9**

*Figure S9. Benchmarking of PM6 QM/MM umbrella sampling force constants* **10**

*Figure S10. PMF dependence on QM region size at the PM6 level.* **11**

*Section S3. Water-mediated stabilization of the pre-reactive APS complex* **11**

*Figure S11. Radial distribution functions (RDFs) of active site waters around the sulfur and phosphorus atoms of APS and AMP.* **12**

*Figure S12. Effect of the [4Fe–4S] cluster redox state on active site organization.* **13**

*Figure S13. Structural analysis of transition state stabilization.* **14**

*Figure S14. Sulfite dipole moment along the reaction coordinate for different redox states of the [4Fe–4S] cluster.* **15**

*Section S4. Local electric field calculations* **15**

*Figure S15. Local electric field along the reaction coordinate for different redox states of the [4Fe–4S] cluster.* **16**

*Figure S16. Per-residue decomposition of the local electric field in the product state.* **16**

*REFERENCES* **17**

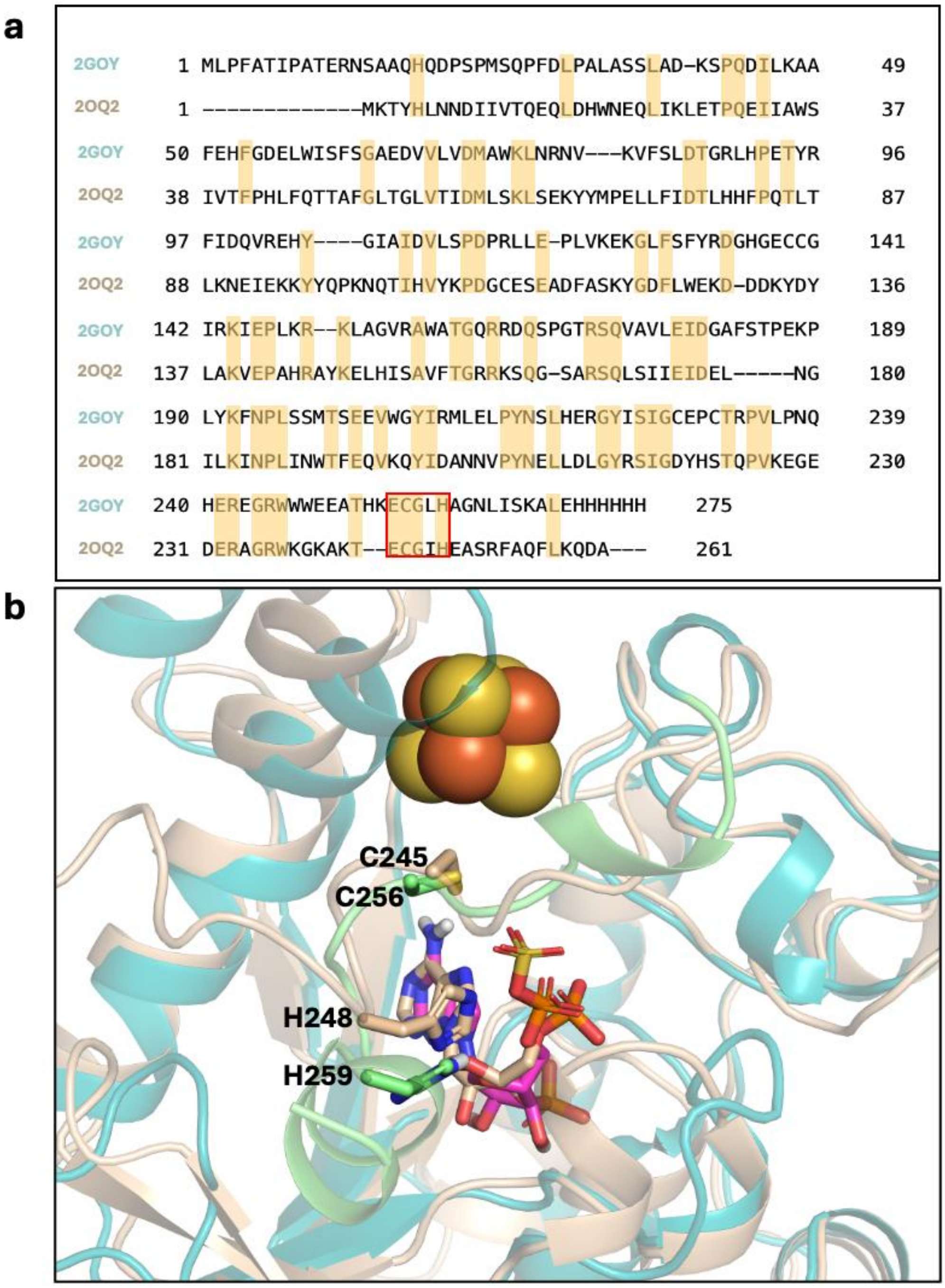


**Figure S1. Sequence alignment and structural comparison of APSR and PAPSR.** **(**a) Pairwise sequence alignment of *Pseudomonas aeruginosa* APS reductase (PDB ID 2GOY) and *Saccharomyces cerevisiae* PAPS reductase (PDB ID 2OQ2), generated with EMBOSS Needle software.[1] The conserved C-terminal ECG(I/L)H motif is highlighted within the red box. (b) Structure superposition of the PAPS reductase crystal structure (sand) with APS reductase (teal), including the AlphaFold-predicted C-terminus (AF-O05927-F1, green). The enzymes bind PAP (sand) and APS (magenta), respectively, revealing analogous active site architectures. The [4Fe-4S] cluster is present in APS reductase but absent in PAPS reductase. Structurally conserved catalytic residues C245/H248 (sand) and modelled C256/H259 (green), shown as sticks, adopt comparable spatial orientations.

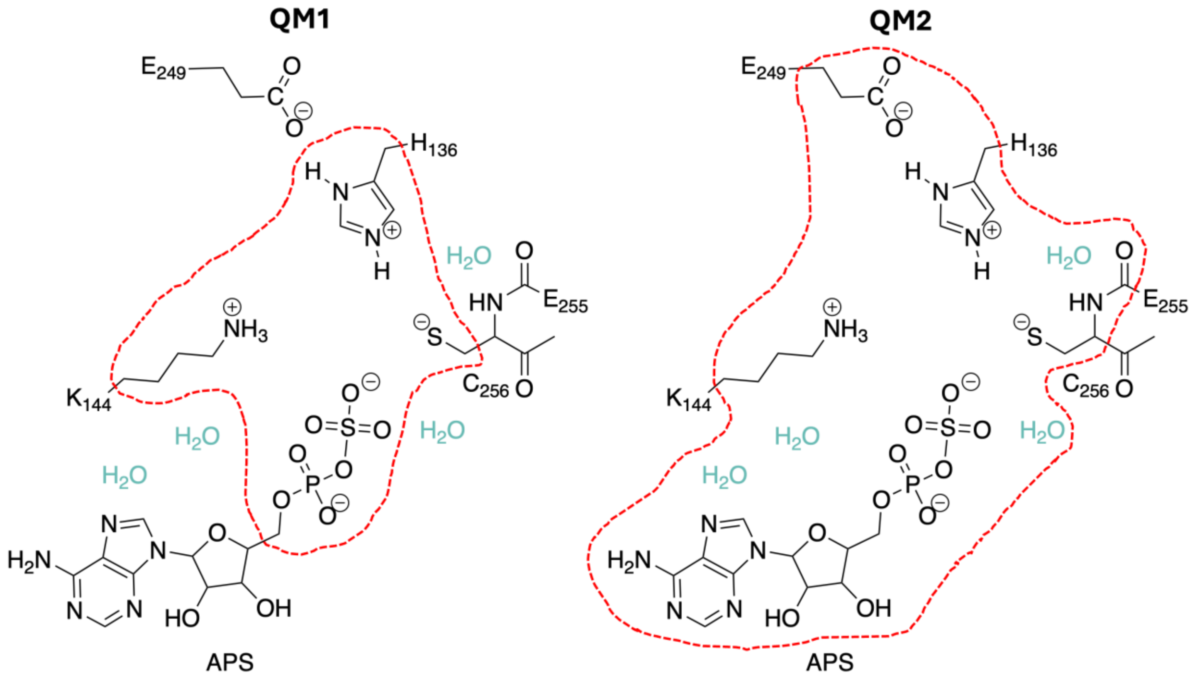


**Figure S2. Definition of QM regions used in QM/MM simulations.** Residues included in the QM regions: QM1 comprises the phosphosulfate moiety of APS plus the Cβ-truncated side chains of H136, K144, and C256 (44 atoms; net charge −1), whereas QM2 expands QM1 to include the full APS substrate, the peptide bond between C256 and E255, the E249 side chain and four water molecules (98 atoms; net charge –2).

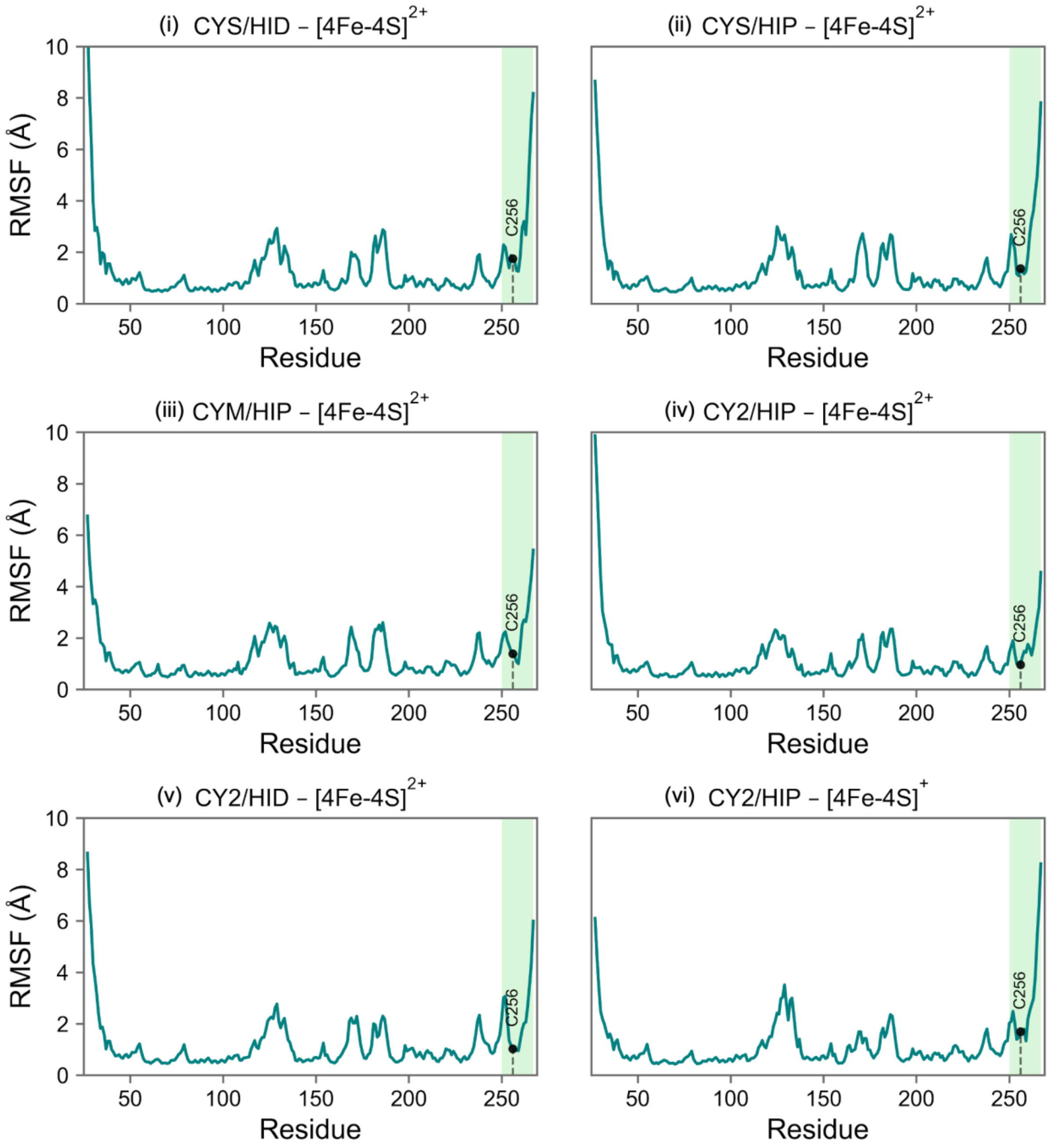


**Figure S3. Comparative protein Cα RMSF analysis of APSR across different protonation C256/H136 and [4Fe–4S] cluster oxidation states.** Protein Cα RMSF profiles reveal changes in residue flexibility, particularly within the C-terminal region, residues 250-267 (highlighted in the green box). The neutral cysteine systems exhibit increased C-terminal flexibility compared with the $[4Fe\text{-}4S]^{2+}$ deprotonated cysteine systems. Relative to CY2/HIP (iv), CY2/HID (v) shows higher C-terminal RMSF, indicating reduced stabilization of this region when H136 is a histidinium, while [4Fe–4S] cluster reduction further promotes its flexibility (vi).

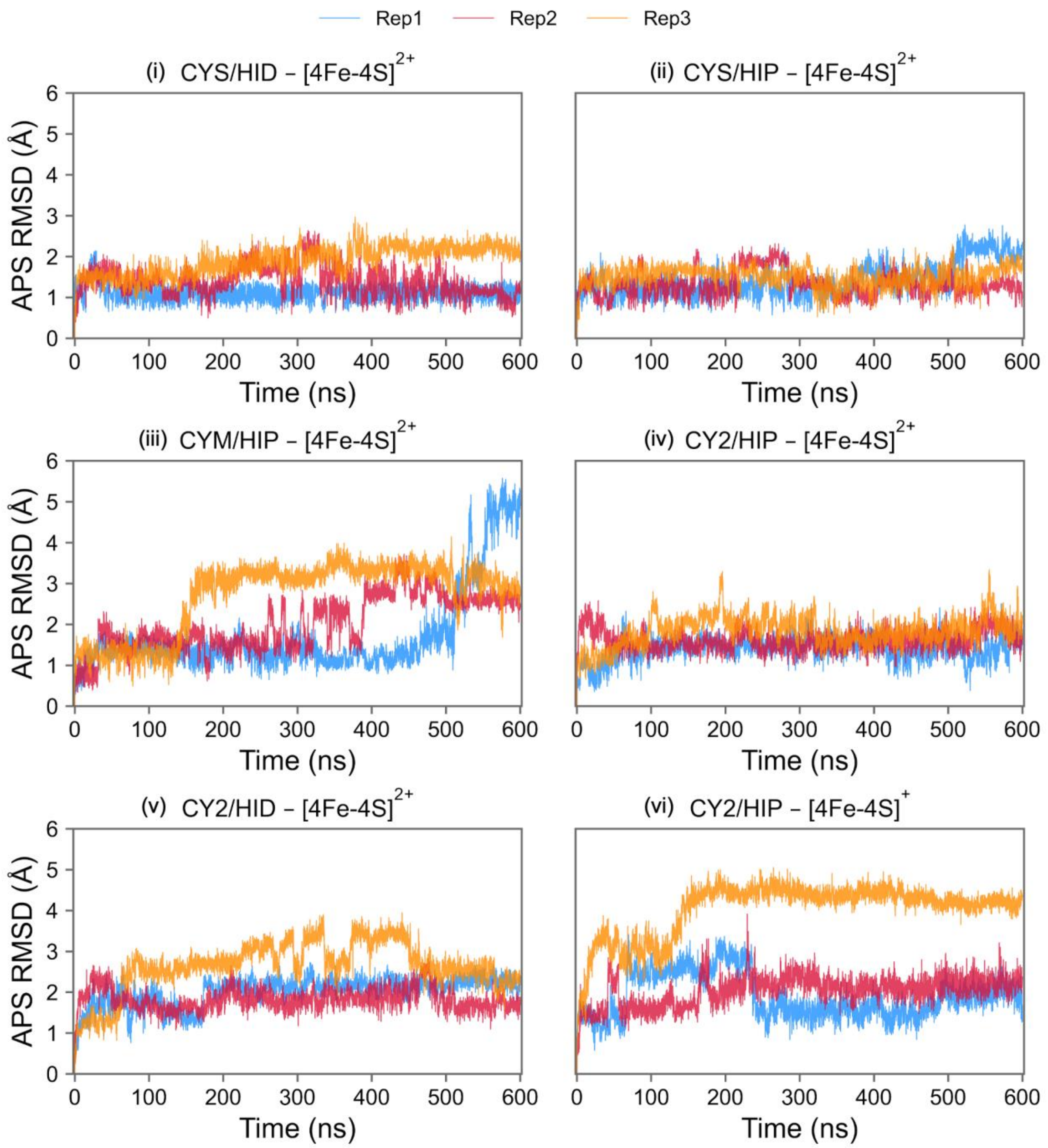


**Figure S4. Comparative APS RMSD analysis of APSR in distinct protonation and [4Fe–4S] cluster oxidation states.** Time evolution APS RMSD was evaluated across three independent replicas per system. The neutral cysteine state (i, ii) maintains a stable ligand binding pose, whereas APS RMSD is moderately increased in CY2/HID (v) compared with CY2/HIP (iv), indicating that HIP protonation better preserves substrate positioning when the catalytic cysteine is deprotonated. Comparison of cluster redox states shows that the oxidized system CY2/HIP (iv) maintains a stable substrate binding pose, with RMSD values generally confined to 1-2 Å, whereas the system with the reduced cluster (vi) displays higher RMSD values and greater inter-replica variability.

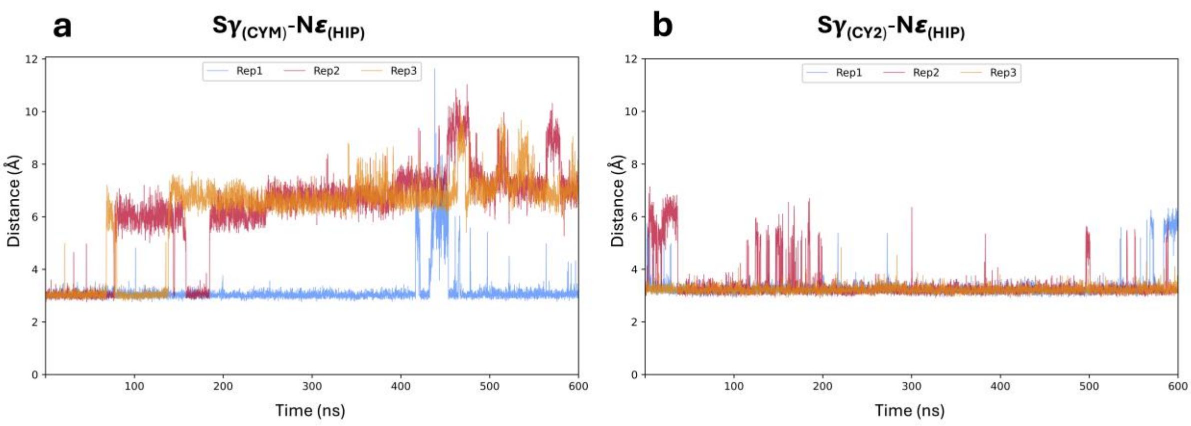


**Figure S5. Thiolate parameters affect active site stability and hydration.** Time evolution of the $S\gamma_{C256}$-$N\varepsilon_{H136}$ distance during MM MD simulations. Default AMBER thiolate parameters (a) yield progressively longer distances with larger fluctuations, indicating weakened hydrogen bonding between C256 and H136, solvent entry into the active site, and consequent substrate destabilization. Custom parameters (b) maintain shorter, more stable Sγ-Nε distances, preserving compact active site geometry, and limiting water accessibility. Mechanistically, the recalibrated terms increase the effective LJ radius of the sulfur atom by 0.15 Å (from the default 2.00 Å in CYM) and shift $S$-$O_{(H_2O)}$ contacts to longer and more realistic distances. This adjustment corrects the default model's short-range oxygen-sulfur contacts and thereby supports a more physically accurate hydration environment.

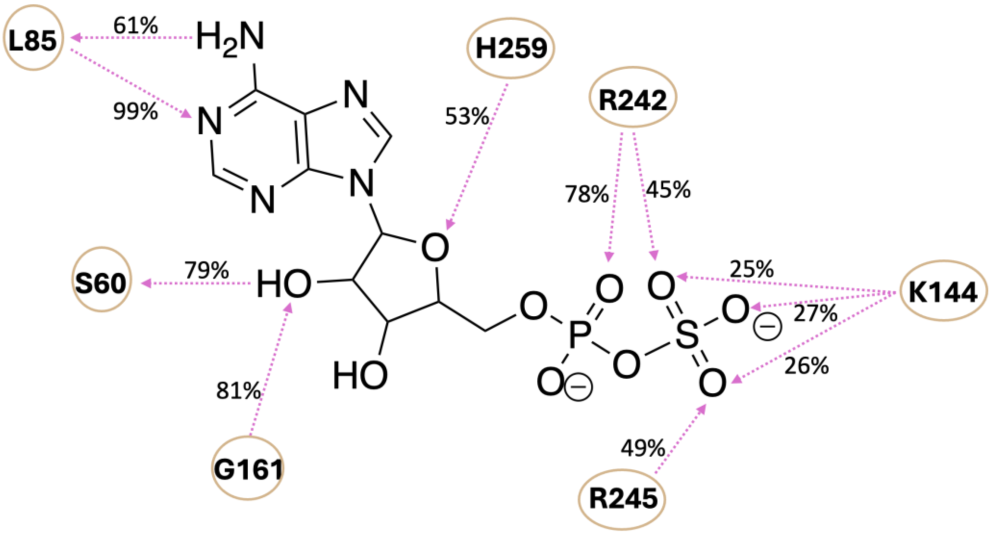


**Figure S6. APSR-substrate hydrogen bond interaction frequency in the CY2/HIP oxidized iron–sulfur cluster system.** Hydrogen bond persistence was quantified from three independent 600 ns MD replicas as the percentage of frames satisfying a donor–acceptor distance threshold of 3.5 Å. Pink dashed arrow indicates the hydrogen-bond direction from donor to acceptor. The adenosine moiety established stable hydrogen bond interactions with S60, L85, G161, H259, while the negatively charged phosphosulfate group is stabilized by K144, R242 and R245.

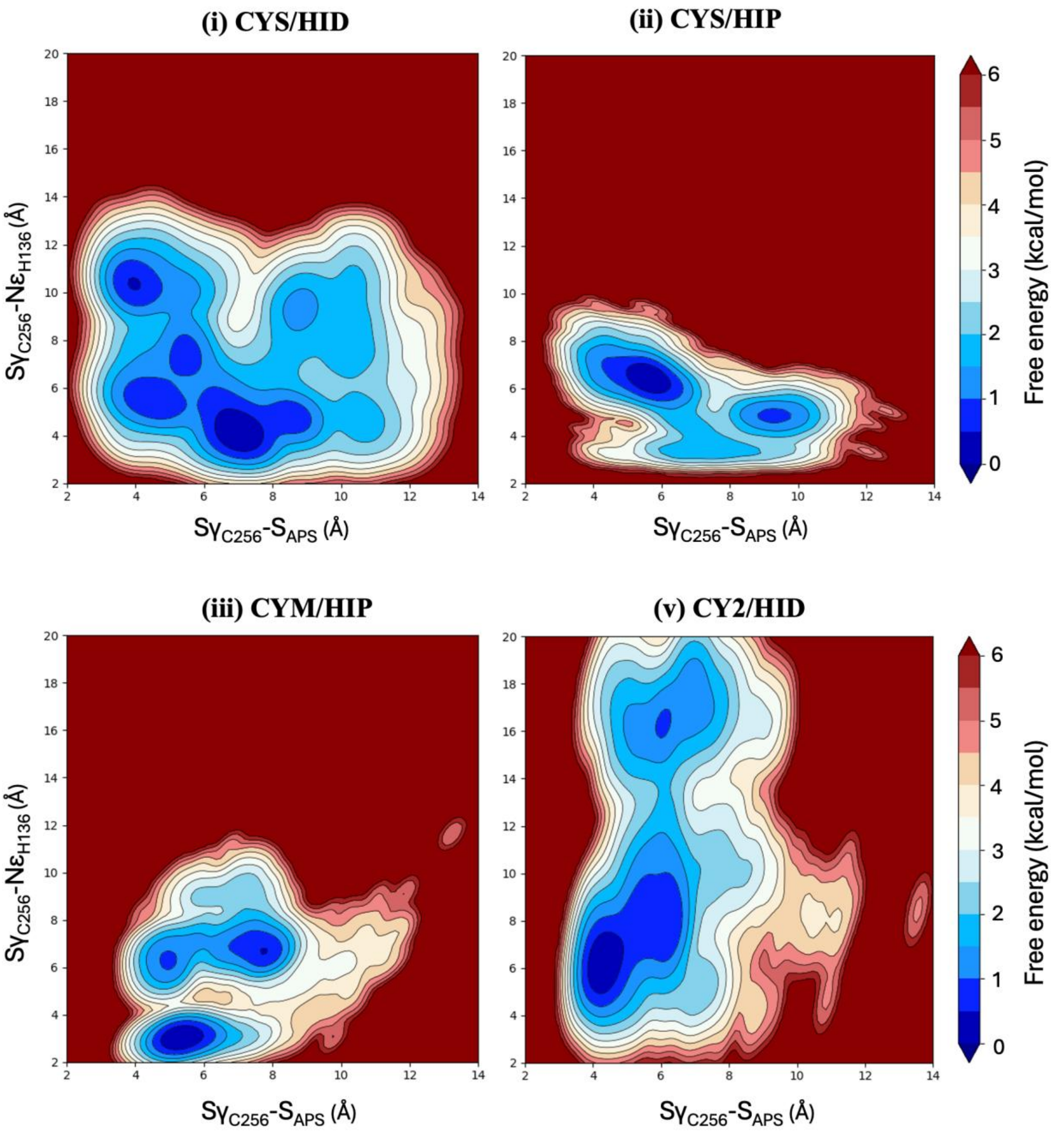


**Figure S7. Free energy landscapes of APSR C256/H136 protonation states.** Free energy landscapes (FELs) of three APSR-substrate systems with distinct C256/H136 protonation state combinations, constructed from MM MD trajectories using the distances $d_1 = S\gamma_{C256}\text{-}N\varepsilon_{H136}$ and $d_2 = S\gamma_{C256}\text{-}S_{APS}$. The resulting landscapes quantify how protonation states modulate active site geometry and C-terminal loop flexibility in the APSR-substrate complex.

#### Section S1. Benchmarking of semiempirical QM/MM methods

Semiempirical QM/MM methods were initially evaluated for the oxidized $[4Fe\text{-}4S]^{2+}$ cluster system to identify a suitable approach for preliminary sampling. The PM6 was selected because it better described the tetrahedral geometry of the phosphosulfate moiety compared with DFTB3 and AM1 (Figure S8).

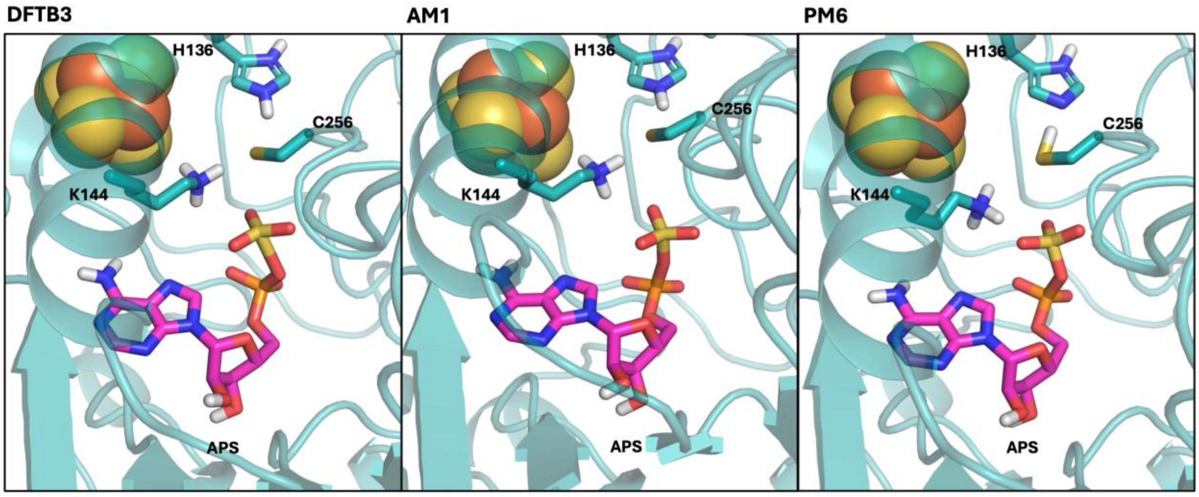


**Figure S8. Semiempirical description of APS phosphosulfate geometry.** Geometries of the phosphosulfate moiety obtained at different semiempirical levels. DFTB3 yields a planar sulfate geometry; AM1 also predicts a planar arrangement with an elongated S-O bond; PM6 produces the most realistic tetrahedral geometry.

#### Section S2. Optimization of the umbrella sampling protocol

Umbrella sampling simulations were initially carried out for the oxidized $[4Fe\text{-}4S]^{2+}$ system using the smaller QM1 region at the PM6/MM level. These preliminary PMF profiles using a force constant of 400 kcal $mol^{-1}$ $Å^{-2}$ yielded high reaction energetics, motivating additional optimization of the umbrella sampling protocol. Accordingly, harmonic force constants of 100, 200 and 300 kcal $mol^{-1}$ $Å^{-2}$ were tested using PM6 with the QM1 region. A force constant of 200 kcal $mol^{-1}$ $Å^{-2}$ was selected because it provided adequate histogram overlap (Figure S9-S10). This optimized protocol was then applied to the larger QM2 region for subsequent QM2 umbrella sampling simulations.

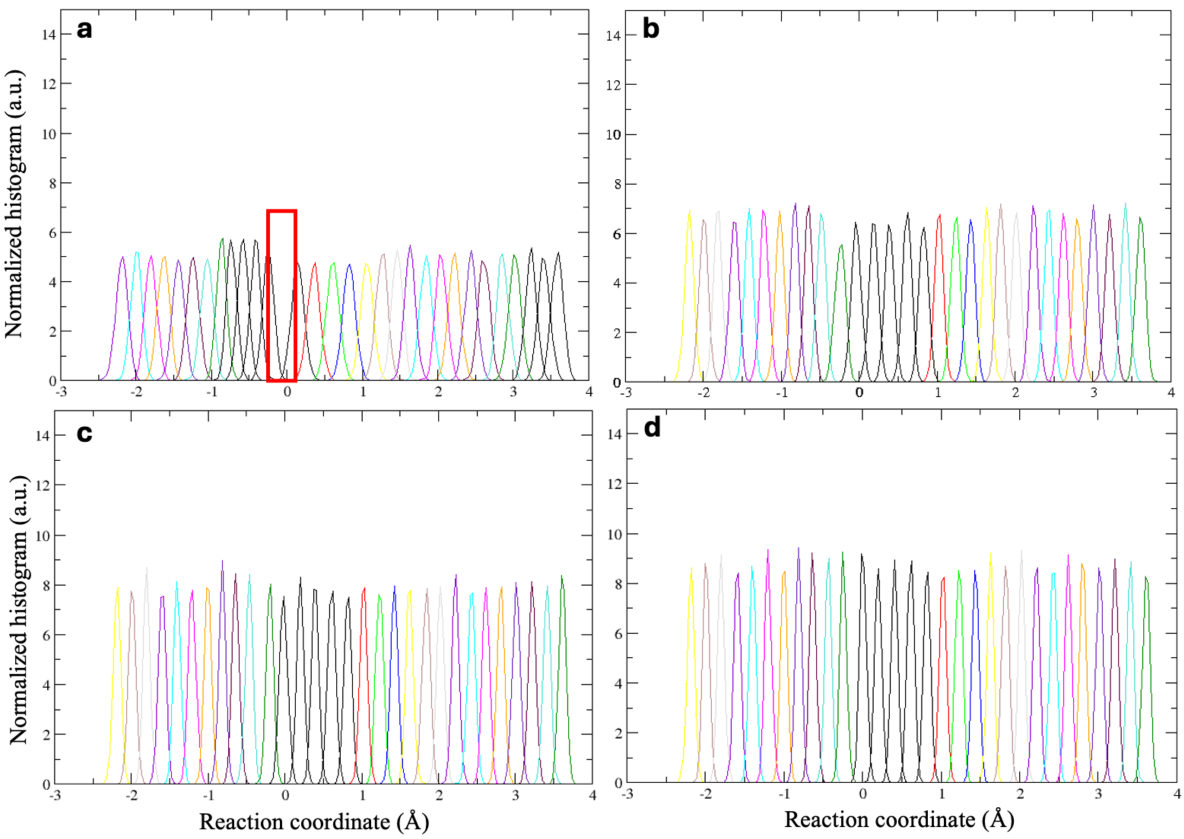


**Figure S9. Benchmarking of PM6 QM/MM umbrella sampling force constants from 100-400 kcal mol$^{-1}$ Å$^{-2}$ and histograms overlap.** Umbrella histograms for force constants of (a) 100, (b) 200, (c) 300, and (d) 400 kcal mol$^{-1}$ Å$^{-2}$. Decreasing the force constant from 400 to 200 kcal mol$^{-1}$ Å$^{-2}$ improved window overlap, whereas a further decrease to 100 kcal mol$^{-1}$ Å$^{-2}$ resulted in insufficient sampling overlap in the pre-transition-state region, highlighted in red. Therefore, a force constant of 200 kcal mol$^{-1}$ Å$^{-2}$ was selected as the optimal value, providing an adequate histogram overlap.

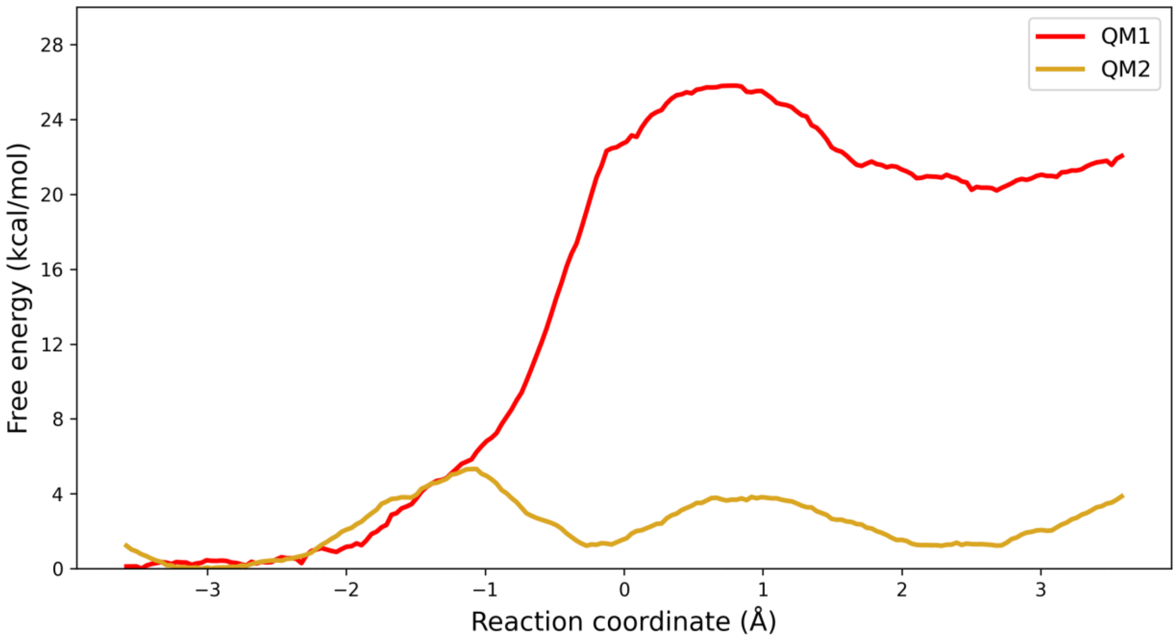


**Figure S10. PMF dependence on QM region size at the PM6 level.** Potentials of mean force (PMFs) were computed using QM/MM MD umbrella sampling simulations with the small QM region (QM1) and the expanded QM region (QM2) at the PM6 semiempirical level.

**Section S3. Water-mediated stabilization of the pre-reactive APS complex**

To better assess the dynamics of the APS reactant and AMP product states, free PM6 QM/MM MD simulations were carried out for both oxidized and reduced [4Fe-4S] cluster systems, using three independent 100 ps replicas for each state. These simulations revealed persistent first hydration shell around the reaction center. Water-phosphate radial distribution function (RDF) analysis revealed differences depending on the iron-sulfur cluster's oxidation state. In the product state, when in presence of the oxidized form of the cluster, RDF showed a sharper and more intense peak near 4 Å, indicating a more persistent water coordination around the phosphate oxygens and a more ordered local solvation shell (Figure S11). This evidence suggests that water molecules populate the cavity generated after covalent attachment of the sulfite group to C256. The higher negative charge of the reduced $[4Fe–4S(Cys)_4]^{3-}$ cluster perturbs active site electrostatics and hydrogen bonding, leading to a more labile hydration shell and broader RDF distributions. In addition, spontaneous proton transfer from H136 to C256 was frequently observed during the unbiased PM6 QM/MM MD simulations, therefore the H136 Nε-H distance was restrained in subsequent umbrella sampling simulations to prevent premature proton transfer and isolate the energetics of the nucleophilic attack step.

Based on these observations, the QM region of the oxidized cluster system was expanded from QM1 to QM2 (Figure S2), comprising the full substrate and four explicit first shell water molecules, to better describe local solvation and polarization effects. Inclusion of these waters in the QM region provided a refined description of the active site and improved stabilization of the substrate through a directional hydrogen bonding network. The water mediated polarization attenuates electrostatic repulsion between the C256 thiolate and phosphosulfate moiety, allowing C256 Sγ to retain sufficient negative character to facilitate the nucleophilic attack.

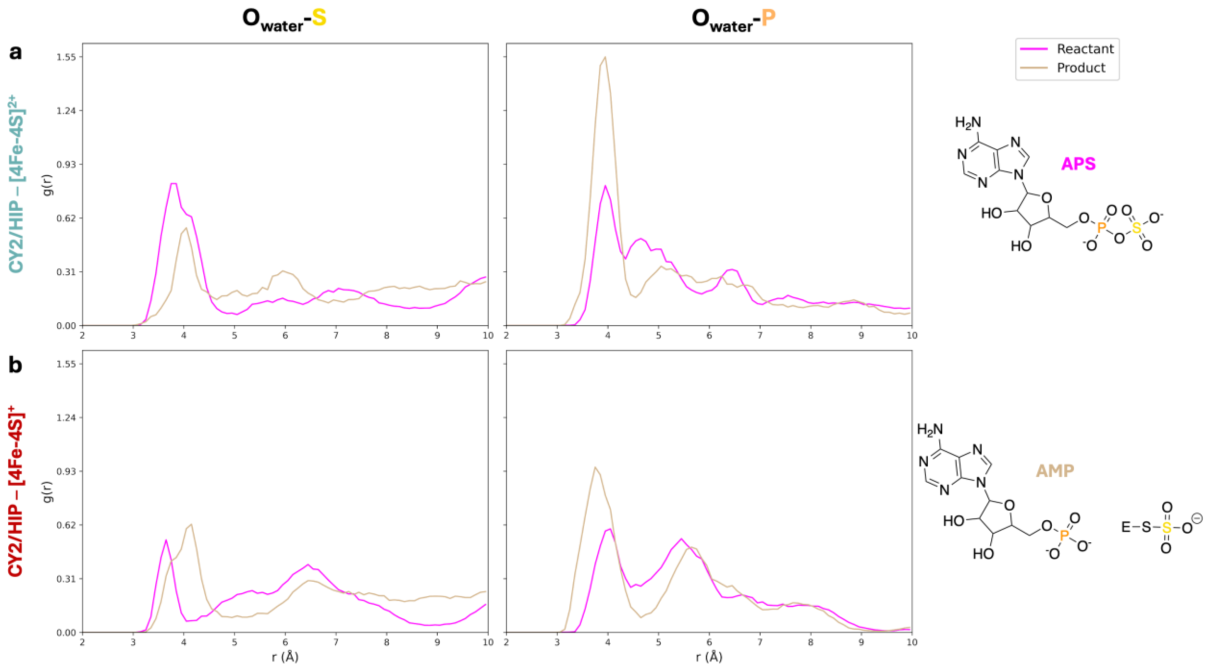


**Figure S11. Radial distribution functions (RDFs) of active site waters around the sulfur and phosphorus atoms of APS and AMP.** RDFs are shown for CY2/HIP systems containing the oxidized $[4Fe\text{-}4S]^{2+}$ cluster (a) and reduced $[4Fe\text{-}4S]^{+}$ cluster (b). Left panels show water oxygen distributions around sulfur atom, whereas right panels show distributions around the phosphorus. In the oxidized product state, the sharper water-phosphate peak near 4 Å reflects more persistent water coordination to the AMP phosphate oxygens. In the reduced system, the broader, lower-intensity distributions suggest faster solvent exchange and a less organized hydration environment.

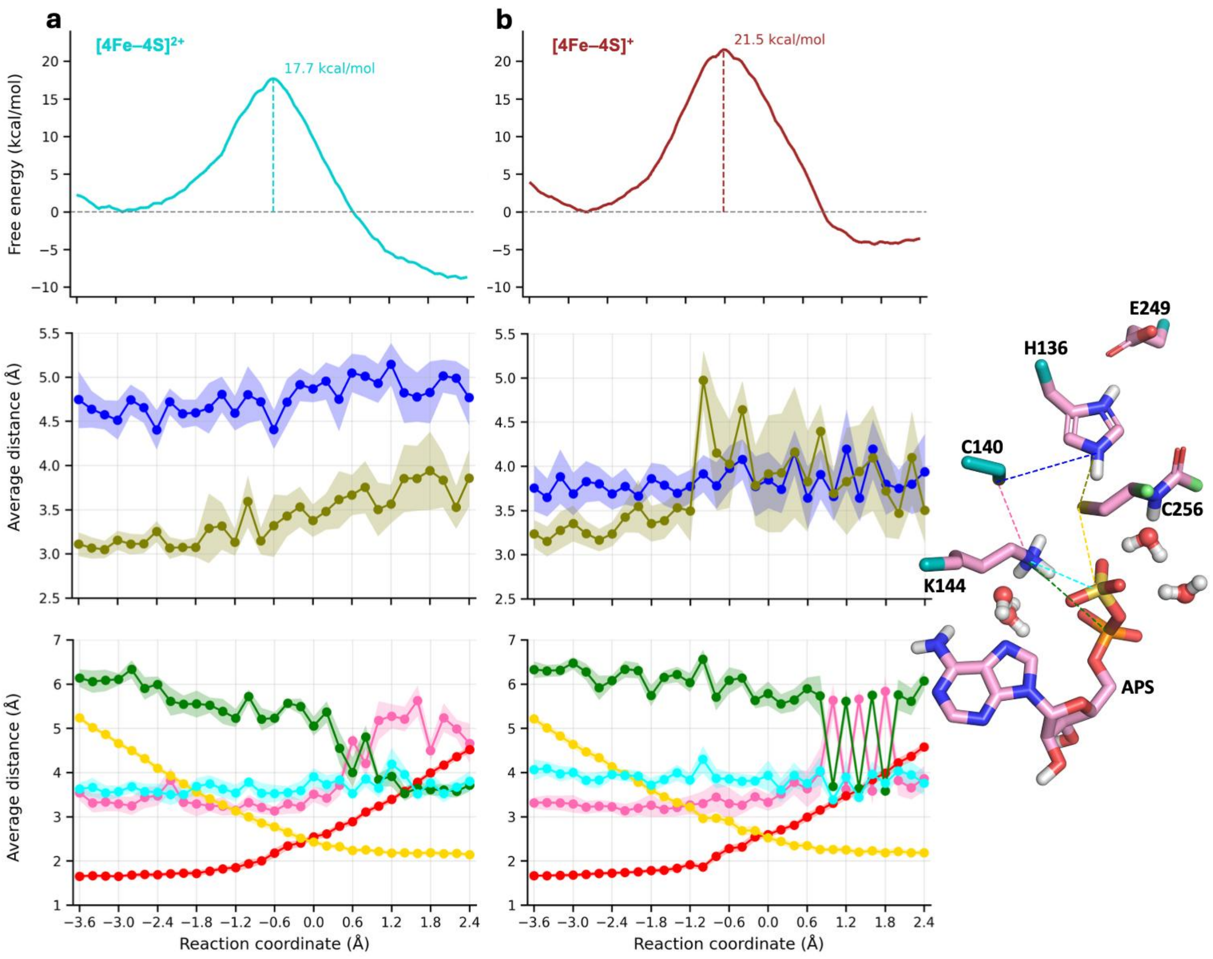


**Figure S12. Effect of the [4Fe–4S] cluster redox state on active site organization.** DFT QM/MM PMFs and average geometric descriptors for the oxidized $[4Fe–4S]^{2+}$ cluster system (a) and the reduced $[4Fe–4S]^{+}$ cluster system (b) along the reaction coordinate. The top panels show the PMF profiles, indicating that cluster reduction increases the activation barrier from 17.7 to 21.5 kcal mol$^{-1}$. The middle and bottom panels report average key distances with standard deviations. The middle panels track the H136 Hε distances to the C140 and C256 cysteine sulfur atoms, shown in blue and olive, respectively. In the reduced system, the overall −3 charge of the cluster favors a closer interaction of H136 with the C140 residue coordinating the cluster, while increasing its separation from catalytic C256 Sγ, thereby reducing electrostatic stabilization of the catalytic cysteine. The bottom panels show distances from K144 N to S_C140 (pink), S_APS (cyan), and P_APS (green), together with the forming C256 Sγ–S_APS bond (yellow) and the breaking S_APS–O_APS bond (red). In the oxidized state, K144 adopts an intermediate position between C140 and APS, maintaining shorter and more stable substrate contacts. By contrast, in the reduced state, K144 remains closer to C140 and farther from APS, with more fluctuating distances to both the product phosphorus atom and the cluster.

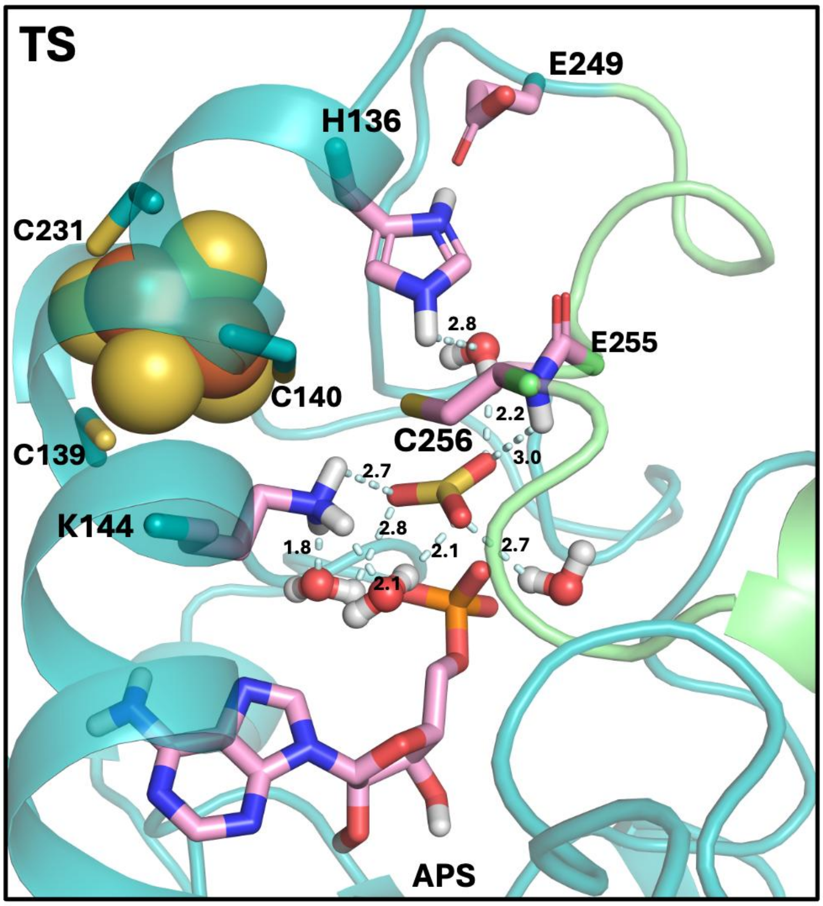


**Figure S13. Structural analysis of transition state stabilization.** Representative snapshot of the transition state, reaction coordinate value of –0.6 Å, showing hydrogen bonds between planar sulfite with K144 and four water molecules (pale cyan dashed lines; distances in Å). Three waters are oriented toward the substrate side of sulfite, while a fourth is stabilized between H136 and C256; the adjacent C256-E255 peptide bond also contributes to transition state stabilization.

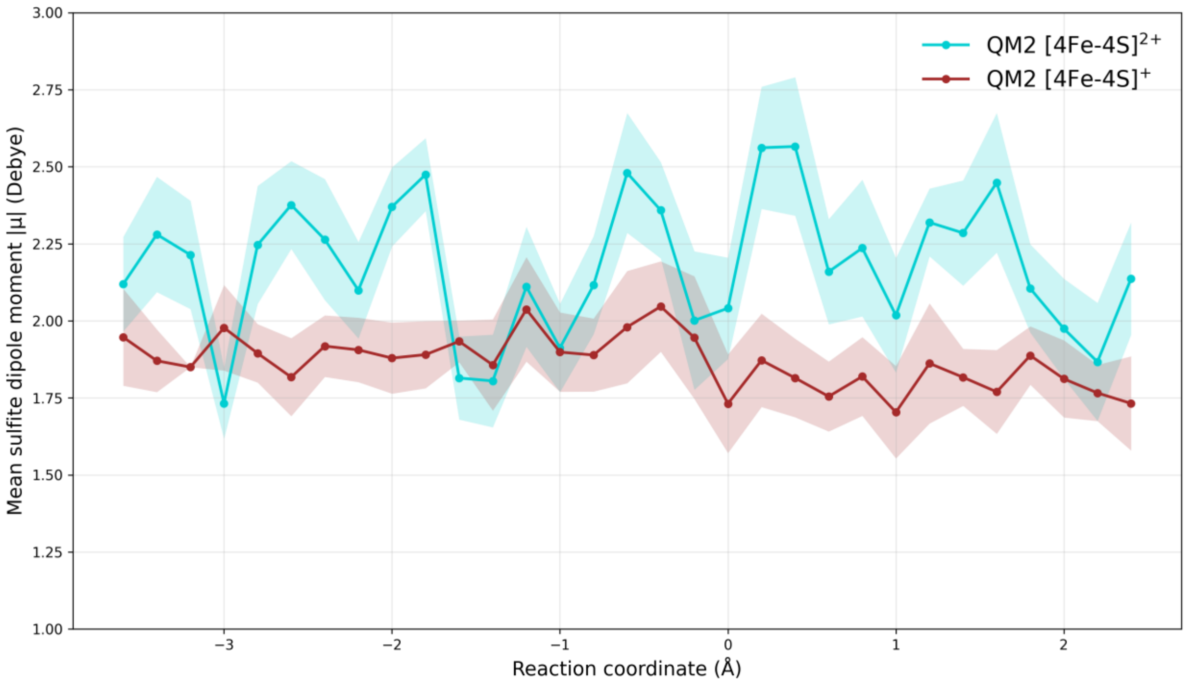


**Figure S14. Sulfite dipole moment along the reaction coordinate for different redox states of the [4Fe–4S] cluster.** Redox-dependent differences within QM2 region suggest that the oxidized system promotes stronger sulfite polarization through the closer K144 side chain, which generates a localized positive electrostatic field toward one sulfite oxygen; this contribution is attenuated in the reduced state due to the larger K144-sulfite separation.

#### Section S4. Local electric field calculations

Local electric fields (LEFs) were calculated along the QM/MM umbrella sampling trajectories using TUPÃ,[2] a Python-based tool that evaluates the instantaneous electric field at user-defined probe points from classical point charges in the simulation. For the APS substrate, the probe was defined as a single oxygen atom located between the sulfate and phosphate groups. The ATOM mode was used to compute the total electric field vector and its magnitude at this atomic coordinate. LEFs were evaluated for 20 equally spaced snapshots taken from the final 2 ps of the QM/MM MD trajectories for the reactant, transition state, and product windows of the umbrella sampling free energy simulations for both the oxidized $[4Fe–4S]^{2+}$ and reduced $[4Fe–4S]^{+}$ cluster states. In each case, the field at the probe oxygen was computed using all protein residues and the iron-sulfur cluster as source atoms, while water molecules, counterions, and the APS substrate itself were excluded, consistent with standard practice for assessing the enzyme-generated field experienced by the reacting moiety. The QM region in the underlying MD was treated at the B3LYP-D3(BJ)/6-31G(d) level, and the resulting electric field values were averaged over the 20 snapshots for each state to characterize the local electrostatic environment along the reaction coordinate.

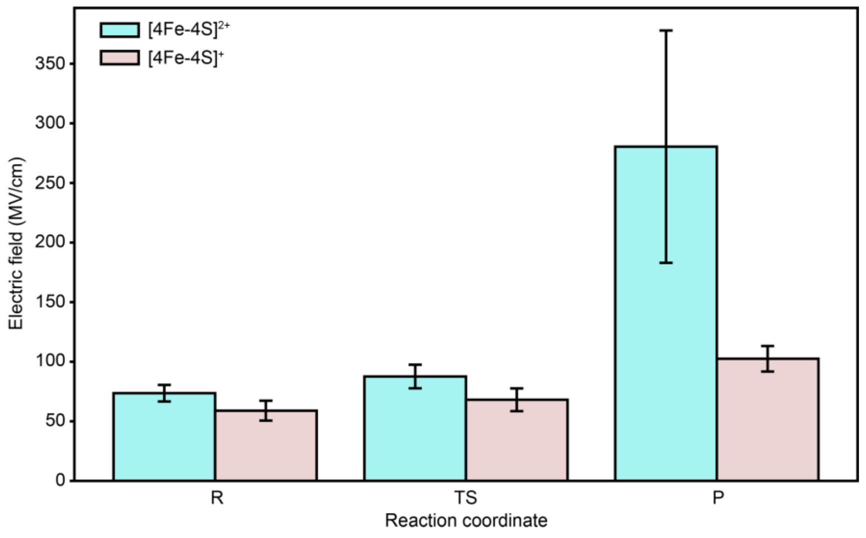


**Figure S15. Local electric field along the reaction coordinate for different redox states of the [4Fe–4S] cluster.** Electric field magnitude at the substrate oxygen along the reaction coordinate for oxidized (cyan) and reduced (pink) [4Fe-4S] cluster states. Bar heights represent average electric field values (MV/cm) with error bars showing standard deviation across the trajectory for reactant (R), transition state (TS), and product (P) intermediates.

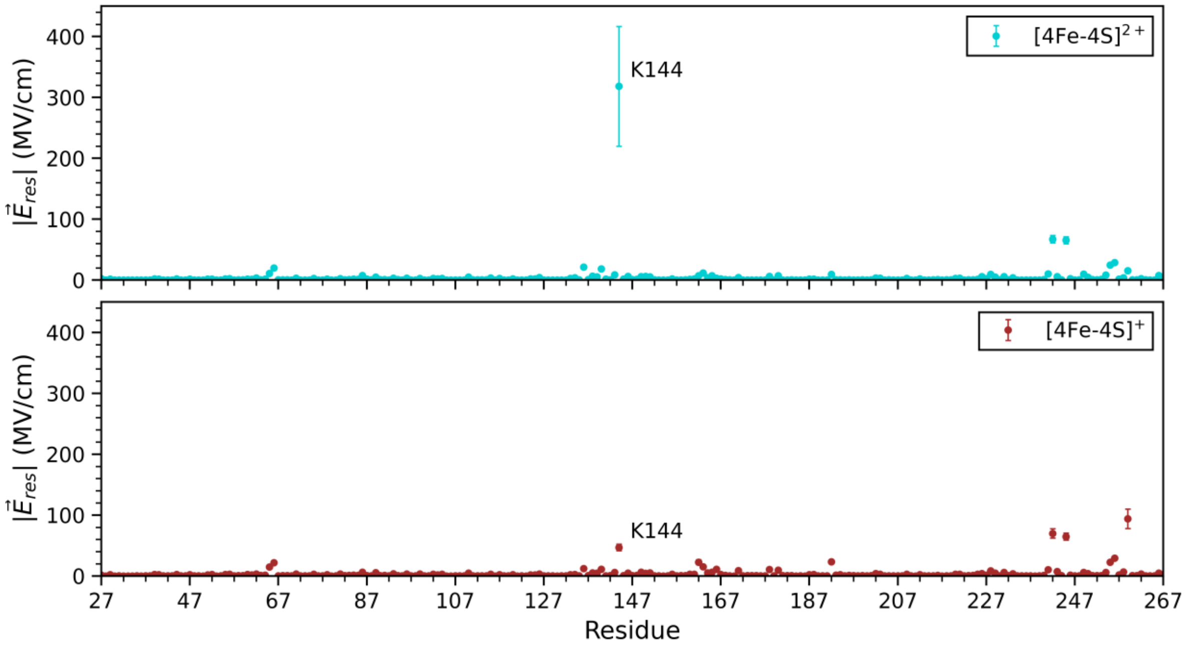


**Figure S16. Per-residue decomposition of the local electric field in the product state.** K144 is the major contributor to the redox-dependent difference in the local electric field, suggesting that changes in the [4Fe–4S] cluster redox state modulate active-site electrostatics predominantly through this residue.

#### REFERENCES


(1) Rice, P.; Longden, I.; Bleasby, A. EMBOSS: The European Molecular Biology Open Software Suite. *Trends in Genetics* **2000**, *16* (6), 276–277. https://doi.org/10.1016/S0168-9525(00)02024-2.

(2) Polêto, M. D.; Lemkul, J. A. TUPÃ: Electric Field Analyses for Molecular Simulations. *J. Comput. Chem.* **2022**, *43* (16), 1113–1119. https://doi.org/10.1002/jcc.26873.
